# Human coronavirus HKU1 neutralization by glycan receptor mimicry

**DOI:** 10.1101/2025.11.05.686863

**Authors:** Ziqi Feng, Nurgun Kose, Fernando R. Moreira, Anne L. M. Kimpel, Jeffrey Copps, Naveenchandra Suryadevara, Daniel J. Jackson, James E. Gern, Ian A. Wilson, Sheng Li, Robert P. de Vries, Ralph S. Baric, Sandhya Bangaru, James E. Crowe, Andrew B. Ward

**Author notes:** These authors contributed equally to this work (Ziqi Feng, Nurgun Kose, Sandhya Bangaru).

## Abstract

Entry of seasonal human coronavirus HKU1 (HCoV-HKU1) into host cells is facilitated by sequential binding to sialoglycans and transmembrane serine protease 2 (TMPRSS2) receptors. However, the neutralizing capacity of antibodies disrupting these receptor interactions have not been examined. Here, we describe the isolation and characterization of a human monoclonal antibody (mAb) HKU1-2 that recognizes the HCoV-HKU1 spike protein and exhibits dose-dependent neutralization of the virus. Epitope mapping and structural analysis revealed that HKU1-2 mAb targets the sialoglycan binding site in the N-terminal domain of the spike protein. A cryo-electron microscopy (cryo-EM) structure of the spike-Fab complex further demonstrated the ability of HKU1-2 to mimic sialic acid binding thereby effectively blocking sialoglycan receptor engagement. HKU1-2 binding is primarily mediated by CDRH3 recognition of NTD residues K80 and W89 that are known to be critical for sialic acid engagement. Overall, our results demonstrate antibody recognition and neutralization of HCoV-HKU1 by receptor mimicry.

## Introduction

Coronaviruses (CoV) are enveloped positive-sense single-stranded RNA viruses and known for their zoonotic threat as evidenced by the epidemic and pandemic outbreaks caused by severe acute respiratory syndrome CoV (SARS-CoV), SARS-CoV-2, and Middle East respiratory syndrome CoV (MERS-CoV) ^1,2^. In addition, four seasonal CoVs, HCoV-229E and HCoV-NL63 of alpha-CoV genus, HCoV-OC43 and HCoV-HKU1 of beta-CoV genus, are currently endemic in the human population ^3,4^. HCoV-HKU1 and HCoV-OC43 belong to the Embecovirus subgenus, and cause mild respiratory infections, with significant health impact in the young, elderly, and immunocompromised populations ^5,6^.

CoV spike (S) glycoprotein on the viral surface is the key protein for host cell attachment and entry and a major target for neutralizing antibodies ^7^. The S proteins of beta-CoVs, which include SARS-CoV, SARS-CoV-2, MERS-CoV, HCoV-HKU1, and HCoV-OC43, are homotrimers consisting of two subunits: S1 contains the amino (N)-terminal domain (NTD) and a carboxy (C)-terminal domain (CTD) that serves as the receptor binding domains, and S2 comprises the fusion machinery required for entry. Spike proteins from diverse CoVs can engage a wide range of receptors and employ distinct entry mechanisms that define the host specificity, tropism and pathogenicity of these viruses ^8^. The spike S1 CTD binds to protein receptors angiotensin-converting enzyme 2 (ACE2) for SARS-CoV and SARS-CoV-2, and dipeptidyl peptidase 4 (DPP4) for MERS-CoV ^3^. The S1 CTDs can exist in various conformational states including the “up” or “open” conformations required for receptor engagement, while the “down” or “close” states are inaccessible ^9-11^. The NTDs, on the other hand, contain an evolutionarily conserved galectin fold with specificity for sialo-glycans. The NTD of MERS-CoV can bind nonmodified α2,3-linked N-acetylneuraminic acid ^12^, while HCoV-HKU1, HCoV-OC43, and bovine-CoV recognize 9-*O*-acetylated sialic acids ^13-18^. A notable exception is SARS-CoV-2, in which both the NTD and CTD have been shown to recognize sialic acids ^19,20^ ^21^.

Frequently, CoVs use both sialoglycan and proteinaceous receptors in tandem to facilitate entry into host cells, in a mechanism previously described for MERS-CoV and SARS-CoV-2 ^20,22^. While HCoV-HKU1 and HCoV-OC43 have been thought to mainly employ their glycan receptor for entry, recent studies have revealed that HCoV-HKU1 also uses its CTD to bind TMPRSS2, a type I transmembrane protein, as its key protein receptor ^23-26^. Furthermore, cryo-EM structures of HCoV-HKU1 spike with sialic acid have shown that sialic acid binding on NTD triggers the HCoV-HKU1 spike CTD into a more open conformation that is accessible for TMPRSS2 binding, suggesting that the binding of the two receptors is sequential with NTD-sialic acid binding being the first key step for entry ^27,28^. Although recent advances have shed light on the mechanisms of HCoV-HKU1 receptor recognition and entry, little is known about whether the HCoV-HKU1 sialoglycan-binding site serves as a target for human antibodies ^28,29^. Furthermore, it remains unclear if antibodies that block initial sialoglycan engagement can effectively inhibit HCoV-HKU1 viral entry.

Here, we isolated a panel of HKU1-reactive mAbs from a healthy donor sample collected prior to COVID-19 pandemic and characterized them based on their binding affinities, neutralization potencies, and epitope specificities. Notably, mAb HKU1-2, which was shown to recognize the NTD of the spike by negative stain electron microscopy (ns-EM), exhibited potent neutralization against the HCoV-HKU1 virus, while mAb HKU1-11, targeting the CTD, showed less neutralization activity. To uncover the molecular mechanisms of engagement and neutralization, we determined high-resolution structures of the HCoV-HKU1 spike in complex with HKU1-2 or HKU1-11 Fabs by single particle cryo-EM. HKU1-11 targeted the CTD at a site distinct from the TMPRSS2 binding site, engaging also in interactions with the N454 glycan. In contrast, HKU1-2 targets an epitope that overlaps with the binding site of 9-*O*-acetylated sialic acid. This interaction was primarily mediated by the long heavy chain CDR3 loop that formed extensive contacts with HCoV-HKU1 NTD. Moreover, the spike NTD epitope residues critical for HKU1-2 binding, as confirmed by HDX-MS and alanine-scanning mutagenesis, were consistent with interactions facilitating sialic acid binding. Functional analysis of HKU1-2 by an HCoV-HKU1 NTD-based hemagglutination inhibition assay demonstrated strong inhibition of 9-*O*-acetylated sialic acid binding to the NTD in the presence of HKU1-2. Overall, our findings uncover sialic acid receptor mimicry as a potent neutralizing mechanism for human CoV antibodies and advance our knowledge of HCoV-HKU1 entry checkpoints that can be targeted for vaccine design and therapeutics.

## Results

### Isolation and characterization of mAbs targeting the HCoV-HKU1 spike

In our previous study, we evaluated human donors with unknown infection histories for the presence of serum (collected pre-COVID-19 pandemic) antibodies to HCoV-HKU1 and HCoV-OC43 spike proteins using electron microscopy based polyclonal epitope mapping (EMPEM) analysis ^30^. Donor 1412, whose serum had the highest HCoV-HKU1 spike-reactive antibody binding titers and displayed multiple antibody specificities, was selected for mAb isolation. The HCoV-HKU1 spike protein used in this study is a soluble spike trimer ectodomain (HKU1-C, isolate N5) containing the 2 proline (2P) stabilizing mutations at residues 1067 and 1068 and the S1/S2 cleavage site mutated to 752-GGSGS-756 ^31^. Hybridoma generation from HCoV-HKU1 spike-reactive B cells led to isolation of a panel of 8 mAbs with half maximal effective concentration (EC50) for binding ranging 10 to 130 ng/mL. Two mAbs, designated HKU1-25 and HKU1-34, also exhibited cross-reactivity to the HCoV-OC43 spike protein, with corresponding EC50 values of 91 and 4,467 ng/mL (**Figure 1A, S1A and S1B**).

**Figure 1.**
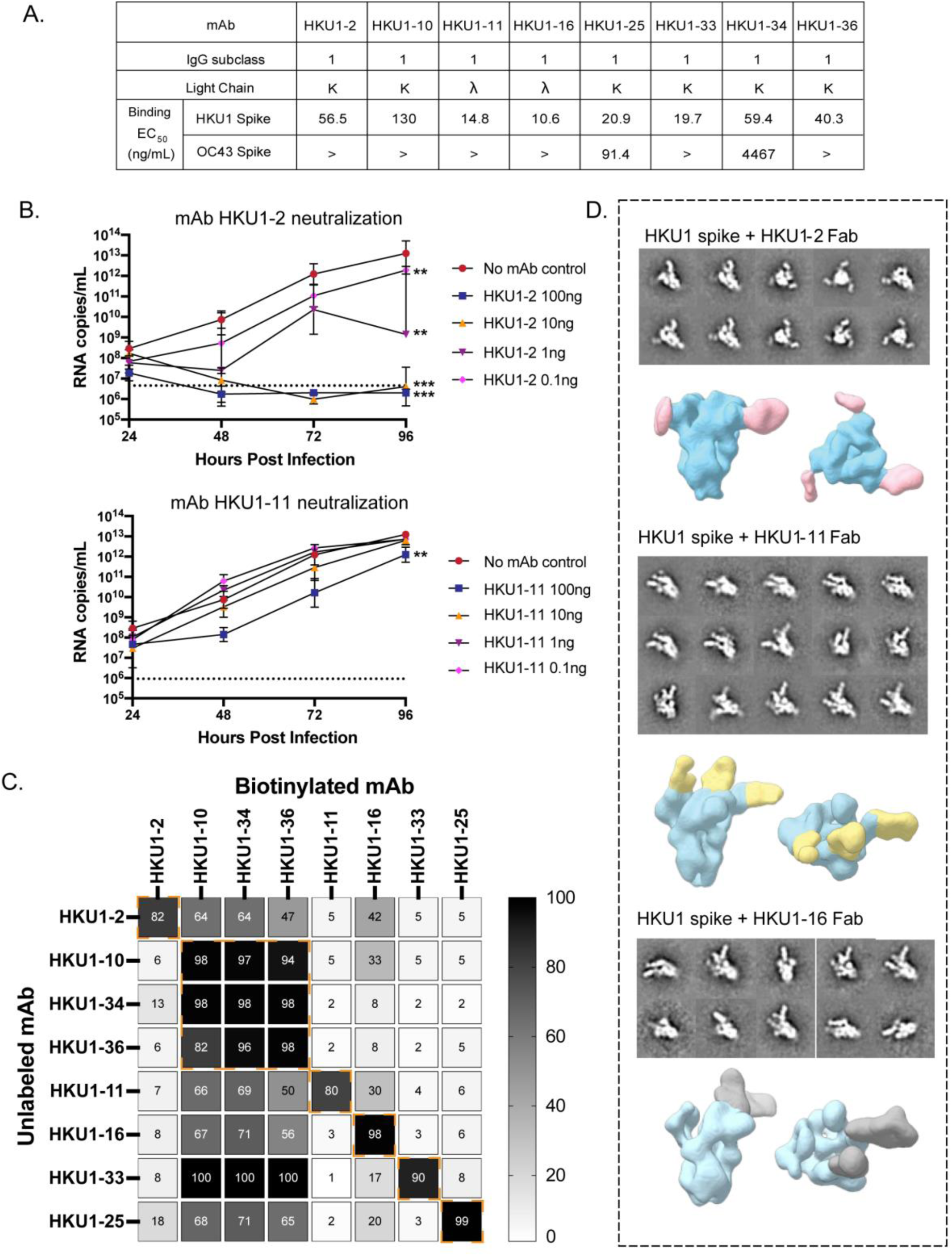
Binding, function and epitope characterization of HKU1-reactive mAbs. (**A**) The IgG subclass, light chain usage and ELISA binding half-maximal effective concentrations (EC50) to spike proteins derived from HCoVs, HCoV-HKU1 and HCoV-OC43, for each of the eight HKU1-reactive mAbs. The > symbol indicates that binding was not observed at concentrations ≤10 µg/mL. (**B**) HCoV-HKU1 viral growth inhibition mediated by mAbs HKU1-2 and HKU1-11. Replication kinetics for HCoV-HKU1 virus was assessed by real-time RT-PCR of RNA isolated from apical washes from HKU1-infected human ciliated airway epithelial cells pretreated with either no mAb (red), or with mAb at concentrations of 100 ng (blue), 10 ng (yellow), 1 ng (violet) or 0.1 ng (magenta) per mL. Titers are represented by virus RNA copies per milliliter. Significant differences in HCoV-HKU1 titers were observed in HKU1-2 treated cells at all concentrations and at 100 ng treatment with HKU1-11. n = 2 for each treatment group. \*\*\**P* < 0.001, \*\**P* < 0.01 (two-way ANOVA test). (**C**) Competition binding results for the panel of 8 HKU1-reactive mAbs as measure by a competition ELISA. The binding signal of the biotinylated reference antibody, measured in the presence of an unlabeled test antibody, was expressed as a percentage relative to the reference antibody binding alone, after background subtraction. The % inhibition is represented by a white-grey scale. Dashed orange boxes indicate different clusters. Only symmetric competition pairs were considered in the clustering analysis. (**D**) Representative 2D class averages and 3D reconstructions from ns-EM analysis of the HCoV-HKU1 spike complexed with either HKU1-2, HKU1-11, or HKU1-16 Fabs. The HCoV-HKU1 spike protein is colored in cyan. Fab HKU1-2, HKU1-11, and HKU1-16 are colored in pink, yellow, and grey, respectively. Both top view and side view are displayed.

To determine the neutralizing capacity of these mAbs, we performed a growth curve analysis of HCoV-HKU1 virus (Genotype B isolate derived from nasal lavage samples) in differentiated polarized cultures of human ciliated airway epithelial cells. The mAbs were tested at concentrations of 0.1, 1, 10, and 100 ng/mL, and viral titers were measured by RT-PCR of RNA extracted from cell supernatants at 24, 48, 72, and 96 hours following treatment with virus-only or virus-mAb mixtures (**Figure 1B and S1C**). Treatment with plasma diluted 10-fold was performed as a negative control. Only HKU1-2 mAb exhibited neutralizing activity, showing dose-dependent viral inhibition, in comparison to other mAbs (**Figure 1B and S1C**). Significant reduction in HCoV-HKU1 titers were observed at lower mAb concentrations of 0.1 and 1 ng/mL, while higher concentrations of 10 and 100 ng/mL reduced titers to below the limit of detection (LOD) (**Figure 1B**). Aside from mAb HKU1-2, the only other antibody to mediate a slight reduction in viral titers [at the highest concentration tested (100 ng/mL)] was mAb HKU1-11 (**Figure 1B**).

We next sought to identify the epitopes targeted by these mAbs by competition binning and ns-EM. Competition binding assays for the binning of eight HCoV-HKU1 mAbs into competition groups of mAbs recognizing common antigenic sites was carried out by sequential binding of an unlabeled mAb followed by a biotinylated mAb on HCoV-HKU1 spike protein-coated plates. The mAbs fell into roughly six competition bins with some cross-competition observed between bins. While HKU1-2 mAb fell into its own competition group, partial inhibition of HKU1-2 binding was observed in the presence of mAbs HKU1-10, -34 and -36 from bin 2 and mAb HKU1-16 (**Figure 1C**). Ns-EM of HCoV-HKU1 spike-Fab complexes revealed that HKU1-11 and HKU1-16 recognize the CTD of the spike, with antibody binding observed across various conformational states of the CTD (**Figure 1D**). Notably, the potent neutralizing mAb HKU1-2 recognized the NTD (**Figure 1D**).

### Structural basis of mAb HKU1-11 binding to the HCoV-HKU1 spike CTD

Based on neutralization data and ns-EM structures, HKU1-2 and HKU1-11 exhibited some neutralizing activity against the HCoV-HKU1 virus and targeted different epitopes. Therefore, we selected these two antibodies for subsequent high-resolution cryo-EM structural studies to elucidate their potential neutralization mechanisms. To generate a better understanding of the epitope targeted by HKU1-11, we performed single particle cryo-EM of the HCoV-HKU1 spike complexed with HKU1-11 Fab at a 1:3 ratio (Fab: spike protomer = 1:1), resulting in a structure at 3.1 Å resolution. We observed HKU1-11 binding to all CTDs on the spike, with defined densities for Fabs bound to the two CTDs in the down conformation, while the Fab density on the CTD in the up conformation was not well resolved due to flexibility (**Figure 2A and S2**). To obtain better resolution at the Fab-spike interface, we determined the local structure of HKU1-11 Fab variable domain with HCoV-HKU1 spike CTD and the adjacent NTD at 3.1 Å resolution (**Figure 2B, S2, and Table S1**). HKU1-11 uses multiple complementarity-determining regions (CDRs) (H2, H3, L1, and L2) to form extensive hydrophilic and hydrophobic interactions with the CTD. HKU1-11 heavy chain residues Y50, S54, S56, D100, and Y100b, form H-bonds with HCoV-HKU1 CTD residues, D570, main-chain of Q559, S568, NAG of N454 glycan, and G449, respectively. Additionally, light chain residue G29 H-bonds with R396, Y32 with S456 and D459, and R50 with main-chains of N454 and S456 (**Figure 2C**). Furthermore, several aromatic and aliphatic residues at heavy chain positions F33, Y52, Y53, Y100b, P100c, I100d, L100g, and light chain residue Y32 on HKU1-11 form hydrophobic interactions with HCoV-HKU1 CTD residues F450, L455, F563, F566, P569, F572, W575, and F577 (**Figure 2C**). While HKU-11 targets the CTD, which engages the HCOV-HKU1 protein receptor TMPRSS2, structural superimposition of our HCoV-HKU1 CTD/HKU1-11 atomic model with the HCoV-HKU1 CTD/TMPRSS2 complex (PDB ID: 8VGT) did not reveal any direct overlap between the HKU1-11 epitope and the TMPRSS2 binding site ^24^ (**Figure 2D**). These results are in congruence with the lack of potent neutralizing capability of HKU1-11, though possible steric hindrance of TMPRSS2 receptor-binding by IgG compared to Fab, might explain the marginal inhibition of virus observed for mAb HKU1-11 at 100 ng/mL.

**Figure 2.**
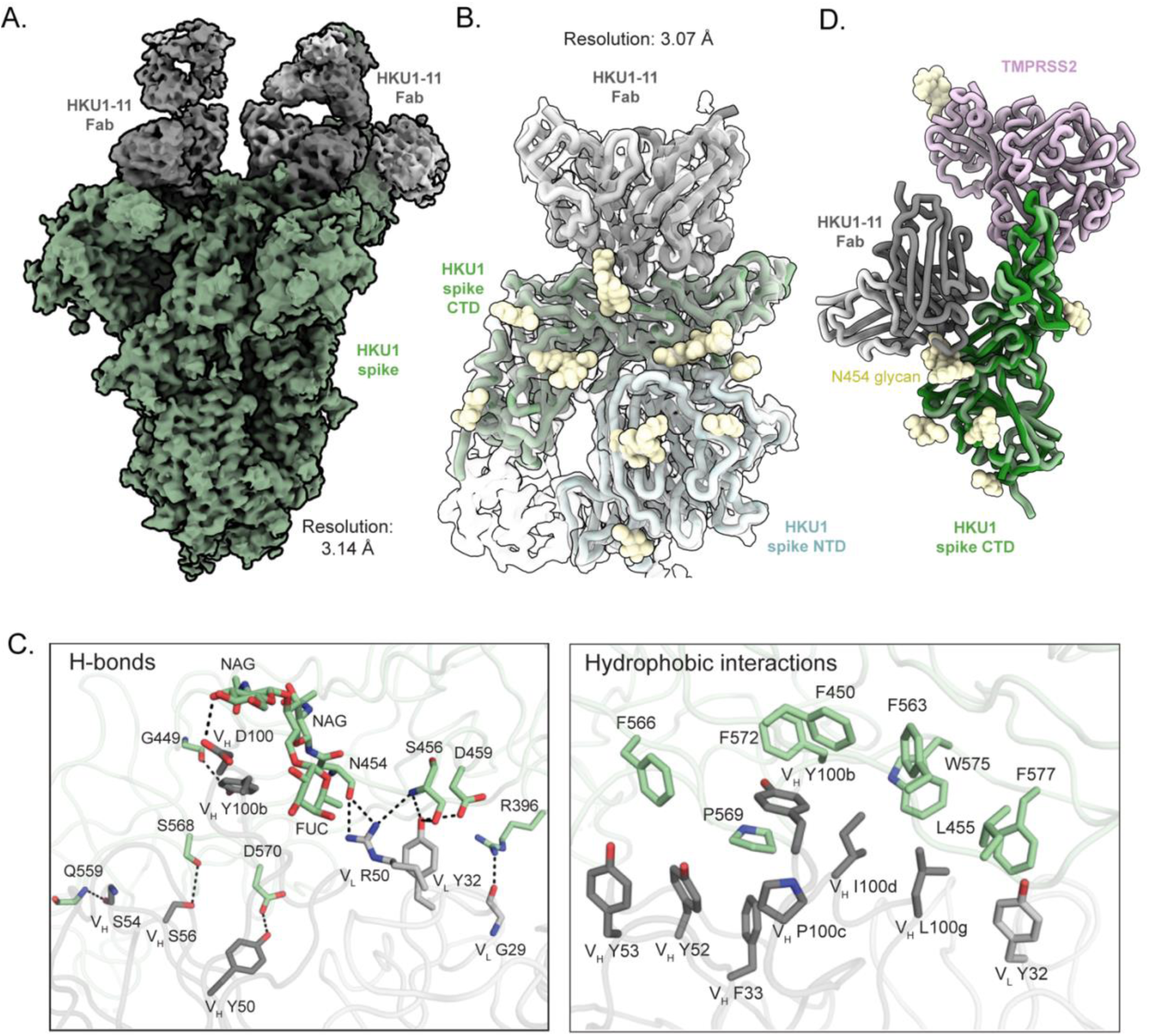
Cryo-EM structure of HCoV-HKU1 spike in complex with mAb HKU1-11. (**A**) Cryo-EM map of HCoV-HKU1 spike trimer with HKU1-11 Fabs bound to the CTD of each protomer, at 3.1 Å resolution. HCoV-HKU1 spike, Fab heavy chain, and light chain are colored in pale green, dark grey, and light grey, respectively. (**B**) Local refine of HCoV-HKU1 spike CTD with HKU1-11 Fab variable domain and the adjacent NTD (3.1 Å resolution) shown as transparent surface. The atomic model built into the local refinement is shown as ribbon representation, with the NTD, CTD, Fab heavy and light chains colored in light cyan, pale green, dark grey, and light grey, respectively. (**C**) Molecular details of the hydrophilic (left) and the hydrophobic (right) interactions between HCoV-HKU1 spike CTD and HKU1-11 Fab. The residues are labeled based on the Kabat numbering scheme. The hydrogen bonds are displayed in the left panel, represented by black dashed lines, while the aromatic and aliphatic residues are displayed in the right panel. (**D**) Superimposition of our HCoV-HKU1 CTD/HKU1-11 Fab complex structure (in this study) with the published HCoV-HKU1 CTD/TMPRSS2 structure (PDB ID: 8VGT) using HCoV-HKU1 CTD as reference, Cα RMSD is 0.8 Å. HCoV-HKU1 CTD and TMPRSS2 are colored in green and pink, respectively. TMPRSS2 does not clash with HKU1-11 Fab.

### Structural insights into the HCoV-HKU1 NTD recognition and neutralization by mAb HKU1-2

Next, we performed cryo-EM analysis of HCoV-HKU1 spike protein complexed with HKU1-2 Fabs to gain structural insight into the molecular mechanisms of HKU1-2 mAb engagement and associated neutralization. We complexed spike with sub-stochiometric concentrations of the HKU1-2 Fab (Fab: spike trimer = 1:1) as attempts with equal molar ratios of Fab per spike protomer led to Fab-induced trimer dissociation. Cryo-EM data processing of the spike-Fab complexes revealed conformational flexibility of the Fab domains as previously observed by ns-EM. These conformations were differentiated by 3D classification with a focus mask around the NTD-Fab region and resolved individually as conformations 1 and 2, both at 3.6 Å resolution (**Figure 3A, 3D, S3, and Table S1**). Structural comparison of atomic models from both states revealed that the spike proteins are nearly identical, with Fabs approaching the NTD at binding angles differing by 5.9° rotating around the Fab axis (**Figure 3B**). HKU1-2, encoded by *IGHV1-24/IGHD2-2*/*IGHJ2* and *IGKV3-11*/*IGKJ4* antibody variable gene segments, interacts with the spike NTD exclusively with its heavy chain. HKU1-2 recognition of the NTD is primarily mediated by its long CDRH3 loop (21 amino acid residues) that is stabilized by an internal disulfide bond between heavy chain residue C98 and C100c, which are both encoded by the *IGHD2-2* gene segment (**Figure 3C**). The CDRH3 loop residues form extensive polar and electrostatic interactions with HCoV-HKU1 NTD: C98, Y100d, and S100f form H-bonds with HCoV-HKU1 NTD N26, S82, and S246, respectively; T99 forms H-bonds with D27 and Y28; and E100e forms salt bridges with K80. In addition to the CDRH3 contacts, E31 on CDRH1 also forms H-bonds with Y84 (**Figure 3F)**. Aromatic and aliphatic residues on CDRH3 (Y100d and P100g) also establish hydrophobic interactions with the NTD (Y28, T31, L79, I83, Y84, and W89) (**Figure 3F**). To facilitate cryo-EM model building, we also determined the crystal structure of HKU1-2 Fab in its apo form at 2.14 Å resolution, with CK mutations (kappa chain constant domain THQGLSSPV to TQGTTSV) added to aid crystallization ^32^ (**Table S2 and Figure S4A**). Structural comparison of apo Fab with spike-bound Fab showed significant differences (high Cα root mean square deviation (RMSD) values) in CDRH2 and CDRH3 loops between structures, highlighting conformational changes that might occur upon NTD engagement (**Figure 3E**).

**Figure 3.**
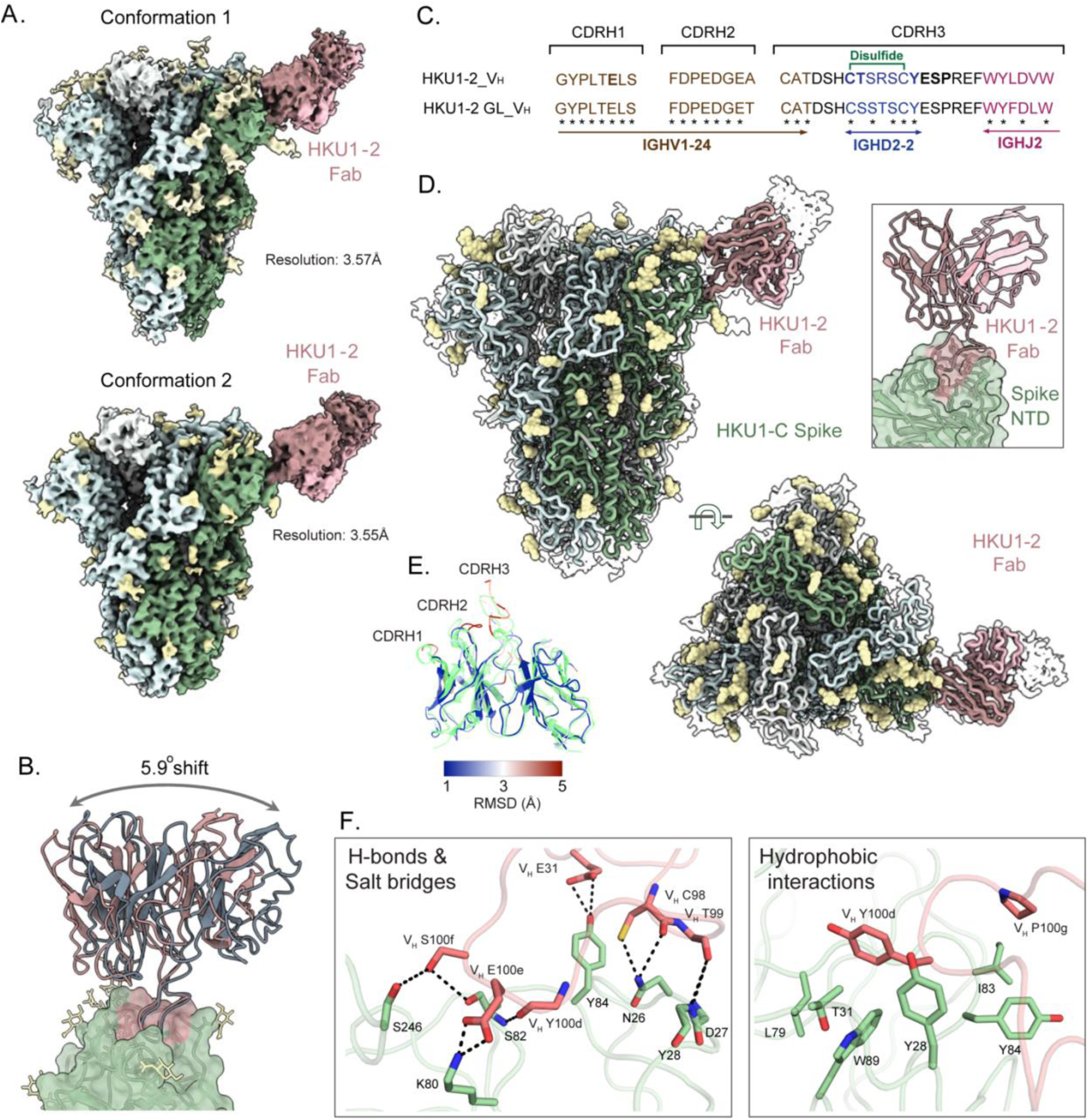
Cryo-EM structure of HCoV-HKU1 spike in complex with mAb HKU1-2. (**A**) Side views of cryo-EM maps of spike-HKU1-2 Fab complex resolved in two different conformational states, both at 3.6 Å resolution. Maps are colored by chain, with the three spike protomers represented in light grey, light cyan, and pale green, the Fab heavy and light chains colored in dark salmon and light pink, and the glycans represented by pale yellow. HKU1-2 Fab binds to the NTD of the prefusion spike, with all CTDs being present in a closed conformation. (**B**) Superimposition of HKU1-2 Fab from the two conformational states (dark salmon and pale blue) show a 5.9-degree shift in their binding angles, calculated using the center of mass for Fab and the NTD. (**C**) IMGT-based sequence alignment of the three CDRH loops between HKU1-2 (VH) and its germline (GL-VH) gene. HKU1-2 heavy chain is encoded by IGHV1-24 (brown), IGHD2-2 (blue), and IGHJ2 (magenta) genes. HKU1-2 residues conserved to its germline are labeled by an asterisk (*), the paratope residues are shown in bold, and the disulfide bond in CDRH3 between C98 and C100c is indicated in green. (**D**) Side and top views of cryo-EM map density (transparent surface) with the docked atomic model represented as a ribbon using the same color scheme as Figure 3A. Zoomed-in view highlights the HKU1-2 heavy chain CDR3 engaging the spike NTD. (**E**) Superimposition of the cryo-EM structure of spike-bound HKU1-2 Fab (colored based on RMSD) and the crystal structure of apo HKU1-2 Fab (transparent green). Structural differences in spike-bound HKU1-2 are color-coded by their RMSDs, and the overall Cα RMSD is 1.2 Å. Dark blue represents low Cα RMSD (< 1Å) and dark red highlights high Cα RMSD (> 5Å). HKU1-2 Fab CDRH2 and CDRH3 have the highest RMSD, highlighting the structural differences between the apo and spike-bound forms. (**F**) Molecular details of the hydrophilic (left) and hydrophobic (right) interactions between the HCoV-HKU1 spike NTD and the Fab. The hydrogen bonds and the salt bridge interactions are displayed in the left panel, represented by black dashed lines, while the aromatic and aliphatic residues are displayed in the right panel.

### HKU1-2 neutralizes HCoV-HKU1 by mimicking and blocking sialic acid receptor binding

Next, we sought to identify the mechanism of HKU1-2 mediated viral growth inhibition. Structural superimposition of atomic models of HCoV-HKU1 CTD-TMPRSS2 and spike-HKU1-2 Fab demonstrated that HKU1-2 does not clash with the protein receptor TMPRSS2, precluding that as a potential neutralization mechanism (**Figure S4B, S4C**) ^28^. While binding to TMPRSS2 is required for HCoV-HKU1 entry into cells, the first key step for initiating the entry process is mediated by spike NTD binding to 9-*O*-acetylated sialic acid receptors ^28^. To investigate if HKU1-2 blocks this initial sialoglycan binding, we superimposed previously published structures of HKU1-C spike NTD in complex with 9-*O*-acetylated sialic acid (PDB ID: 8Y8J) and HKU1-A NTD with GD3 ganglioside (PDB ID: 9BTB) with our spike-HKU1-2 Fab structure ^28,33^ (**Figure 4A**). Remarkably, the CDRH3 of mAb HKU1-2 engages the same pocket as that occupied by 9-*O*-acetylated sialic acid and GD3 ganglioside, forming similar hydrophilic and hydrophobic interactions with the HCoV-HKU1 NTD. Several interactions across 9-*O*-acetylated sialic acid, GD3 ganglioside, and mAb HKU1-2 engagement of the spike NTD were conserved, including H-bonds with residues N26, K80, and S246, and hydrophobic interactions with residues L79, Y84, and W89 (**Figure 4B**). Among the three HCoV-HKU1 genotypes (A, B, and C), spike proteins from genotypes B and C are highly homologous while sharing ∼76% similarity with genotype A^29^. Despite differences, the key NTD residue contacts shared by mAb HKU1-2 and sialic acid binding are conserved across all three genotypes, highlighting their importance in receptor binding and the potential for broad HKU1-2 reactivity (**Figure S5A**). Indeed, HKU1-2 mAb displayed comparable affinities to recombinant spikes from genotypes HKU1-A, HKU1-B and HKU1-C at 12.3 nM, 0.7 nM and 2.7 nM, respectively (**Figure S6**).

**Figure 4.**
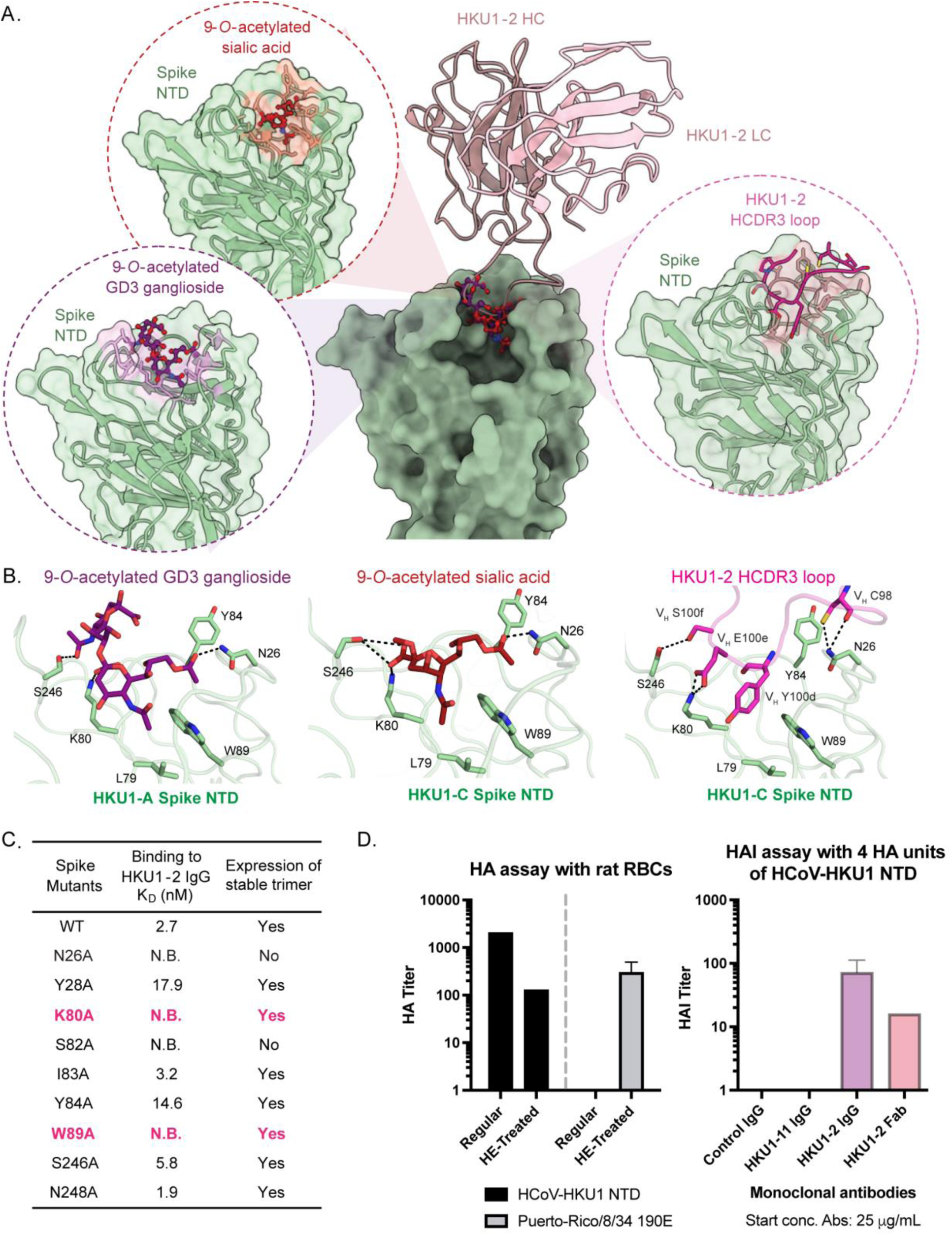
HKU1-2 mimics sialic acid receptor binding to HCoV-HKU1 NTD and blocks the interaction. (**A**) Comparison of HCoV-HKU1 NTD/HKU1-2 Fab complex structure (this study) with the structures of HCoV-HKU1 NTD complexed with 9-*O*-acetylated sialic acid (PDB ID: 8Y8J) and 9-*O*-acetylated GD3 ganglioside (PDB ID: 9BTB), using HCoV-HKU1 NTD for alignment. Cα RMSDs are 0.9 Å and 0.6 Å, respectively. HCoV-HKU1 NTD, HKU1-2 CDRH3 loop, 9-*O*-acetylated sialic acid, and 9-*O*-acetylated GD3 ganglioside are shown in pale green, magenta, brick red, and purple, respectively. Zoomed-in views highlight their engagement of the same binding pocket on NTD. (**B**) Detailed interactions between HCoV-HKU1 spike NTD and 9-*O*-acetylated GD3 ganglioside (left), 9-*O*-acetylated sialic acid (middle), and HKU1-2 Fab CDRH3 (right). Hydrogen bonds are represented by black dashed lines; aromatic and aliphatic residues are displayed. (**C**) Effects of single alanine mutations in the spike on binding to HKU1-2 IgG and on stable trimer expression. Binding affinity is expressed as the nanomolar dissociation constant (*K*D). **(D)** HA of rat RBCs, which have the 9-*O*-acetylated sialic acid receptor (left) and inhibition of NTD-NP mediated HA activity by HKU1-2 (right). The left panel shows the ability of HCoV-HKU1 NTD-NP and PR8 190E hemagglutinin to hemagglutinate rat RBCs that are untreated or HE-treated. HE-treatment removes 9-*O*-acetylated sialic acids on the rat RBCs. The right panel shows the results from an HAI assay using 4 HA units of NTD-NP. Serial dilutions of mAbs, HKU1-2 (IgG and Fab), RSV-specific IgG and HKU1-11 IgG at a starting concentration of 25 µg/mL were tested for HAI activity. The HAI titers, expressed as the highest dilution (lowest concentration) of mAb that inhibits 4 HA units of NTD-NP, demonstrate inhibitory activity for both HKU1-2 IgG and Fab at concentrations of 0.98 µg/mL and 3.13 µg/mL, respectively.

To further validate our finding that HKU1-2 mimics sialic acid receptor engagement, HCoV-HKU1 spikes with single alanine substitutions in the sialic acid binding pocket were generated and tested for binding with HKU1-2 IgG using biolayer interferometry (BLI) (**Figures 4C, S5B and S6**). Among the spike mutants tested, two mutants (N26A and S82A) reduced overall yield and trimer stability, suggesting their necessary role in spike folding/expression. Three mutant spikes (Y28A, Y84A, and S246A) decreased HKU1-2 binding affinity by 6.6-, 5.4-, and 2.1-fold, respectively, compared to WT, while two mutants (K80A and W89A) completely abrogated binding to HKU1-2, a majority of these residues being critical for sialoglycan interactions ^13,27^. Two other mutants (I83A and N248A) had minimal effects on the binding affinity. Overall, our results demonstrate that mAb HKU1-2 engages the spike NTD through a sialic acid receptor mimicry mechanism.

To elucidate if this structural mimicry leads to functional blocking of sialic acid receptor binding, we performed hemagglutination inhibition (HAI) assay using rat red blood cells (RBCs), which display the 9-*O*-acetylated sialic acid receptor for HCoV-HKU1 ^13^. Although the spike NTD-glycan interactions are of low affinity, avidity can be increased via multivalent interactions to facilitate hemagglutination (HA) assays ^12,14,34^. To increase multivalency, we employed a pA-LS nanoparticle (NP) scaffold to display 18 HCoV-HKU1 NTD dimers ^12,35^, which bind efficiently to rat RBCs ^13^. As a control for the presence of 9-*O*-acetylated sialic acid on rat RBCs, we employed an influenza A virus (A/Puerto Rico/8/1934 190E (PR8 190E)) hemagglutinin trimeric protein, precomplexed using α-strep and α-human antibodies ^36^, whose glycan binding is inhibited by presence of 9-*O*-acetylated sialic acid ^37^. The 9-*O*-acetylated sialic acids on the rat RBCs can be removed by pretreating them with bovine coronavirus hemagglutinin-esterase (BCoV HE) ^38^; therefore, 9-*O*-acetylated sialic acid depletion by BCoV HE was expected to increase PR8 190E hemagglutinin binding to rat RBCs. Indeed, HA of rat RBCs by PR8 190E hemagglutinin was only observed for BCoV HE-treated RBCs (**Figure 4D and Figure S7A**), thereby confirming the presence of 9-*O*-acetylated sialic acids on rat RBCs. To verify HCoV-HKU1 NTD binding of 9-*O*-acetylated sialic acid on rat RBCs, we tested the HA activity of the NTD NP on BCoV HE-treated and untreated rat RBCs. While NTD NP-based HA was observed for untreated rat RBCs, treatment with BCoV HE significantly decreased HA of rat RBCs (**Figure 4D and Figure S7A**). To determine whether the HKU1-2 mAb inhibits NTD NP-mediated HA of rat RBCs, we performed an HA inhibition assay with mAb HKU1-2, as both IgG and Fab, along with control IgGs (a respiratory syncytial virus (RSV)-specific mAb and mAb HKU-11). The HKU1-2 mAb in both formats exhibited considerable HAI activity whereas the control IgG showed no detectable HA inhibition (**Figure 4D and Figure S7B**). The HAI titer is expressed as the highest dilution (lowest concentration) of antibody that inhibits HA at four hemagglutination units (4 HAU) of HCoV-HKU1 NTD. The lowest concentrations at which HKU1-2 IgG and Fab exhibited HAI activity was measured at 0.98 µg/mL and 3.13 µg/mL, respectively (**Figure 4D and Figure S7B**). These results indicate that inhibition of NTD-mediated sialic acid receptor engagement is the main mechanism of HKU1-2 neutralization.

### HKU1-2 binding alters HCoV-HKU1 spike dynamics revealed by HDX-MS

Attachment of HCoV-HKU1 spike NTD to 9-*O*-acetylated sialic acid receptors has been observed to facilitate CTD binding to the protein receptor TMPRSS2 by triggering CTD into an “up” conformation ^28^. While HKU1-2 binding to sialic acid binding pocket did not appear to trigger this “up” conformation, we assessed if HKU1-2 induces any changes in spike dynamics using hydrogen-deuterium exchange mass spectrometry (HDX-MS). Calculation of deuteration differences of the spike amide backbone in the presence or absence of HKU1-2 Fab at various incubation times (10, 100, 1,000, and 10,000 s), revealed decreased deuterium exchange at spike residues 79 to 89 in the presence of HKU1-2 Fab, consistent with reduced accessibility to epitope residues (**Figure 5 and S8**) ^27,28,39^. We also observed lower and higher deuterium exchange rates at non-epitope spike segments: 576-577, 583-592, 1161-1164, and 51-53, 650-661, 994-998, respectively. Notably, all of these segments except for 1161-1164, were located at various interprotomer interfaces (**Figure 5 and S8**). Together, these results show that HKU1-2 induces changes in the spike dynamics. Collectively, our study identified and characterized a potently neutralizing mAb HKU1-2, which primarily functions by blocking sialic acid receptor binding and potentially altering dynamics of the spike trimer.

**Figure 5.**
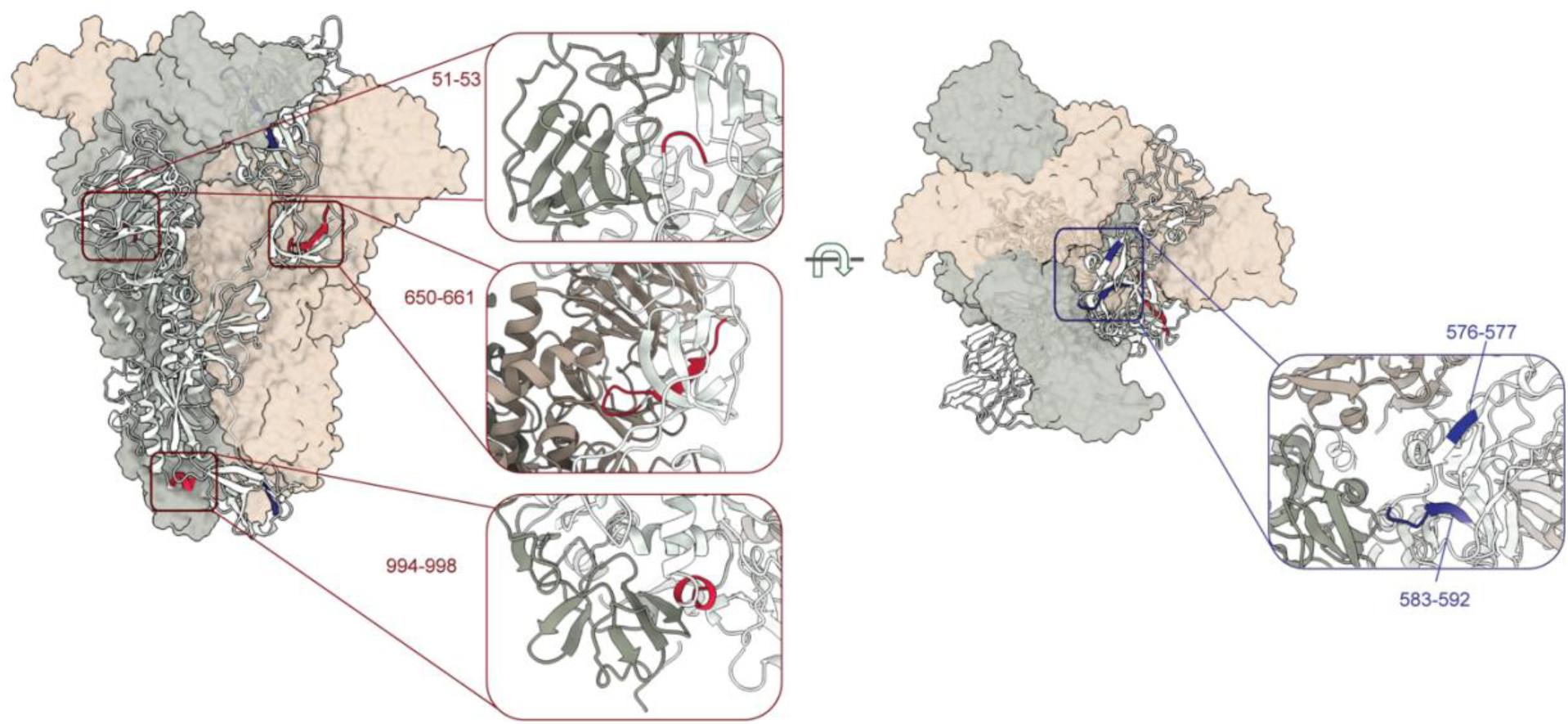
HDX-MS reveals HCoV-HKU1 spike dynamics induced by HKU1-2 mAb. Differences in the deuteration of HKU1-2 mAb-bound spike compared to the apo-spike. The deuteration differences are labelled on a single spike protomer represented as a ribbon, with regions of faster exchange shown in red and those with lower exchange colored in blue. The other protomers shown as semi-transparent surface are colored in pale green and sand. Zoomed-in views highlight the faster and slower deuterium exchange rates upon HKU1-2 binding, occurring at the inter-protomeric interfaces. Their residue numbers are listed. The deuteration differences in the HKU1-2 epitope residues are not displayed here.

## Discussion

Coronaviruses can employ multiple receptors to infect host cells, which offers advantages for efficient viral entry and broader tissue tropism, thereby representing an adaptive strategy for effective transmission. Multi-receptor usage also enhances viral infectivity and host range as evident in human coronaviruses such as SARS-CoV-2 and MERS-CoV. In general, CoV spikes can engage both glycan and protein receptors synergistically to facilitate viral entry ^20,40^. This entry mechanism has been shown to occur in a sequential manner, where initial binding to a sialic acid receptor is often thought to trigger conformational changes in the spike domains. This facilitates binding to protein receptors, which is often coupled with molecular ratcheting mechanisms required for fusion peptide exposure and membrane fusion ^40-42^. Recent studies have shed light on HKU1’s dual usage of the cell receptors, further demonstrating the sequential viral entry mechanism ^24,25,28^. HCoV-HKU1 spike protein NTD attaches the sialoglycan receptor 9-*O*-acetylated sialic acid first, which allosterically stimulates the CTD into an “up” conformation, thereby enabling binding to its protein receptor TMPRSS2. Isolation of neutralizing antibodies capable of blocking these spike-receptor interactions not only offer valuable insight into these underlying molecular mechanisms but also serve as potential candidates for therapeutics and for identifying targets for vaccine design approaches. To this end, we isolated mAbs from a convalescent donor resulting in identification of nine mAbs, which were further characterized based on their binding affinities, neutralizing properties and epitope bins. Most notably, we identified a potently neutralizing mAb HKU1-2 that recognizes the HCoV-HKU1 NTD and some weakly neutralizing mAbs targeting the CTD.

Cryo-EM analysis of a representative CTD mAb, HKU1-11, did not show direct overlap between HKU-11 epitope and the protein receptor TMPRSS2-binding site. However, as an IgG, there is potential for steric hinderance to TMPRSS2 binding, which may explain the weaker neutralization. Our study does not dismiss the neutralizing ability of HCoV-HKU1 CTD-targeting mAbs, as previous studies have identified neutralizing antibodies targeting the CTDs of both HCoV-HKU1 and HCoV-OC43 spikes ^29,43^. Interestingly, structural analysis of the potently neutralizing mAb HKU1-2 by cryo-EM revealed that it employs molecular mimicry to engage the 9-*O*-acetylated sialic acid binding site on the HCoV-HKU1 spike NTD, which was confirmed by loss of mAb binding to spike mutants with alanine substitutions at residues critical for 9-*O*-acetylated sialic acid binding. This mode of engagement translated to functional inhibition of receptor binding, as demonstrated by an hemagglutination inhibition assay using rat RBCs, which display 9-*O*-acetylated sialic acid receptors on their surface. This receptor-mimicking mechanism has been documented in influenza virus, where some antibodies against hemagglutinin or neuraminidase use a long CDRH3 loop to insert into the receptor-binding site and compete with sialic acid through molecular mimicry ^44,45^. Further, HKU1-2 altered dynamics at the inter-protomeric interface, and such binding may inhibit functional molecular ratchet motions required for cell entry and present a dual mechanism of neutralization. Although HCoV-HKU1 infection is generally mild in healthy adults, it can cause severe, life-threatening lower respiratory tract infections, particularly in elderly patients or those with underlying medical conditions, sometimes progressing to acute respiratory distress syndrome (ARDS). These clinical features highlight the potential therapeutic use of the HKU1-2 antibody, as it may help prevent severe disease caused by HKU1 in vulnerable populations.

Spike binding to sialic acid receptors have been demonstrated for several beta-CoVs, including SARS-CoV-2, MERS-CoV, HCoV-OC43 and HCoV-HKU1. Disrupting these spike-sialic acid interactions offers an effective strategy for preventing infection. For SARS-CoV-2, acetylated sialic acid-derived glycoclusters were shown to potently inhibit cell binding and viral infectivity^46^. However, antibody-based inhibition of sialic acid binding has, so far, been reported only for a single HCoV-OC43 spike antibody, 46C12, which was derived from a spike-immunized H2L2 mice ^43^. MAb 46C12 specifically targets the spike NTD and engages the 9-*O*-acetylated sialic acid receptor binding pocket. To our knowledge, HKU1-2 is the only naturally occurring human antibody identified to target this binding site and effectively inhibit receptor binding. This study also provides the first direct evidence for disruption of sialic acid receptor engagement as a mechanism for potent HCoV-HKU1 neutralization, highlighting this site as an effective target for therapeutic strategies.

## Acknowledgements

We thank Hannah L. Turner and Charles A. Bowman for their help with electron microscopy, data acquisition and data processing. We thank Lauren Holden for her assistance with the manuscript. The Wilson lab is grateful to the staff of the National Synchrotron Light Source II (NSLS-II) beamline 17-ID-2 for assistance. This research used resources of NSLS-II, a U.S. Department of Energy (DOE) Office of Science User Facility operated for the DOE Office of Science by Brookhaven National Laboratory under Contract No. DE-SC0012704. We thank Henry Tien for technical support with the crystallization robot. We thank Robyn Stanfield for assistance in data collection and Marc-André Elsliger with computation.

## Author contributions

S.B., Z.F., N.K., J.E.C., and A.B.W. conceptualized the studies. N.K. performed mAb isolation and binding characterization. Z.F., S.B., and J.C. expressed recombinant proteins. D.J.J. and J.E.G. provided human samples. S.B. and Z.F. performed cryo-EM data collection, processed the cryo-EM data and built models. S.B. and Z.F. collected and processed ns-EM data. Z.F. performed binding assays and interpreted data. Z.F. acquired and analyzed X-ray data. F.R.M. performed neutralization assays. N.S. performed competition binning assay. A.L.M.K performed HA and HAI assays. S.L. performed HDX-MS experiments. I.A.W, R.P.dV, R.S.B, S.B, J.E.C, and A.B.W supervised research. Z.F., S.B., J.E.C., and A.B.W. wrote the paper, and all authors reviewed and edited the paper.

## Data and materials availability

All data needed to evaluate the conclusions in the paper are present in the paper and/or the Supplementary Materials. Cryo-EM maps have been deposited at the Electron Microscopy Data Bank (EMDB) with accession codes: EMD-73658 (HKU1 spike-HKU1-11 Fab), EMD-73659 (HKU1 spike-HKU1-2 Fab state1), and EMD-73660 (HKU1 spike-HKU1-2 Fab state2). Atomic models have been deposited to the RCSB Protein Data Bank (PDB) with PDB IDs: 9YYW (HKU1-2 Fab), 9YYX (HKU1 spike-HKU1-11 Fab), 9YYY (HKU1 spike-HKU1-2 Fab state1), and 9YYZ (HKU1 spike-HKU1-2 Fab state2).

## Materials and Methods

### Human samples used in the study

PBMCs were isolated from blood collected from donor 1412. The studies were approved by the Institutional Review Board of Vanderbilt University Medical Center. Samples were obtained after written informed consent.

### Recombinant expression of HCoV-HKU1 and HCoV-OC43 spike ectodomains and alanine mutants

All spike ectodomain constructs contain a C-terminal T4 fibritin trimerization domain, an HRV3C cleavage site, an 8×His-Tag, and a Twin-strep-tag for purification. The HCoV-HKU1 spike construct includes residues 1 to 1276 from isolate N5 (GenBank Q0ZME7) with the S1/S2 cleavage site modified to 752-GGSGS-756 and the residues 1067 to 1068 replaced by prolines for generating stable uncleaved spike proteins. Nine single alanine-substituted variants were generated based on this construct, including N26A, Y28A, K80A, S82A, I83A, Y84A, W89A, S246A, and N248A. These positions were predicted to be possible binding sites for sialic acid and HKU1-2 mAb. The HCoV-OC43 spike construct contains spike residues 1 to 1287 (GenBank AIL49484.1) with introduction of stabilizing prolines at sites 1079 and 1080.

### Expression and purification of recombinant spike proteins

For protein expression, Expi293F cells (Thermo Fisher Scientific: A14527, RRID: CVCL_D615) were transected with the spike plasmid of interest. Transfected cells were incubated at 37°C with shaking at 110 rpm for 7 days for protein expression. The spike proteins were purified from the supernatants with Ni Sepharose excel (Cytiva: 17-3712-03) using a 250 mM imidazole elution buffer, and buffer-exchanged to TBS (20 mM tris and 150 mM NaCl, pH 7.4) before further purification with Superdex 200 16/90 column (GE Healthcare Biosciences). Protein fractions corresponding to the trimeric spike proteins were collected and concentrated. The quality of purified proteins was assessed by ns-EM.

### PBMC isolation and hybridoma generation

Generation of human hybridoma cell lines secreting human mAbs was performed as described previously ^47^. Peripheral blood mononuclear cells (PBMCs) were isolated from a single adult donor (Blood collected pre-COVID-19 pandemic) using the Ficoll-gradient method and cells were frozen and stored at -80°C for later use. For hybridoma generation, cells were thawed and transformed with Epstein Barr virus and plated in 384-well plates to generate immortalized lymphoblastoid cell lines. The plates were incubated for seven days at 37°C and the supernatants were screened for the presence of antibodies reactive to the HCoV-HKU1 spike protein using a capture enzyme linked immunosorbent assay (ELISA). B cells secreting HCoV-HKU1 spike-reactive antibodies were expanded on 96-well plates and electro-fused with HMMA2.5 myeloma cells using an ECM 2001 electro cell manipulator (BTX) to generate stable hybridoma cell lines. The hybridomas were cloned by single-cell flow cytometric sorting into 384-well plates to obtain homogenous antibody secretion. Cells were extended after the clonal selection and submitted for antibody variable gene DNA sequence analysis.

### MAb production and purification

For expression of hybridoma-derived mAbs, clones were grown in Serum Free Media (SFM; Thermo Fisher). Culture supernatants were purified using HiTrap Mab Select Sure (Cytiva) on a 24-column parallel chromatograph system (Protein BioSolutions). Gibco ExpiCHO Expression System using transfected ExpiCHO cell cultures was used for recombinant expression of HKU1-2 IgG. Purified mAbs were routinely tested for endotoxin levels and found to have less than 30 EU/mg IgG. The tests were performed using the PTS20IF cartridge (Charles River) with a sensitive range of 10 to 0.1 EU mL and an Endosafe NexGen-MCS instrument (Charles River).

### Determination of mAb binding affinities as half maximal effective concentrations (EC50)

EC50 concentrations for mAb binding to spike proteins were measured by ELISA. Briefly, 384-well microtiter plates were coated with purified HCoV-HKU1 or HCoV-OC43 spike proteins at 2 μg/mL in carbonate buffer pH 9.7, and incubated overnight at 4°C. Plates were blocked with 5% non-fat dry milk and 2% goat serum in Dulbecco’s phosphate-buffered saline (D-PBS) containing 0.05% Tween-20 (D-PBS-T) for 1hr. For EC50 calculations, the plates were incubated with three-fold serial dilutions of each mAb (each dilution performed in triplicate) starting at 10 µg/mL followed by incubation with 1:4,000 dilution of anti-human IgG alkaline phosphatase conjugate. The plates were washed three times between each step with PBS containing 0.1% Tween-20. The binding of antibodies was detected using goat anti-human IgG conjugated with horseradish peroxide (HRP) (Southern Biotech, 2040-05, 1:4,000 dilution) and TMB substrate (Thermo Fisher). The color development in the wells was monitored, and the reaction was stopped by adding 1M HCI. Absorbance was measured at 450 nm using a spectrophotometer (Bio-Tek). EC50 values were calculated using Prism v.9.0 software (GraphPad) after log transformation of the mAb concentrations using sigmoidal dose-response nonlinear regression analysis.

### HCoV-HKU1 neutralization assay

Human Ciliated Airway Epithelial cells were used for inoculation, recovery, and culturing of HCoV-HKU1 virus from nasal lavage samples. HCoV-HKU1 was cultured from a clinical specimen obtained from a child during an acute infection. Cells were seeded at 2.5E5 cells/mL on permeable Transwell-COL supports and maintained for 4-6 weeks to form differentiated polarized cultures. For inoculation, the apical side of the membrane was washed 3× with 1×-Calcium Magnesium Free PBS (1×-CMF PBS) and inoculated with 200 μL of a 1:1 1×-CMF PBS and nasal wash mixture. Following a 2 hr incubation at 32°C, the inoculum was removed, and the membrane washed for 10 minutes with 500 μL of 1X-CMF PBS. Supernatant was collected by conducting a wash on the apical side for 10 minutes with 120 μL of maintenance media or 1×-CMF PBS at various time points, clarified and frozen at -80°C. RNA from clarified washes was extracted by TRIzol-Zymo kit and the virus yield quantified using One-Step RT-PCR in copies/mL. For virus neutralization assay, the viral growth curve analysis was carried out as described in the presence of mAbs at concentrations of 100 ng, 10 ng, 1 ng, and 0.1 ng per mL. Supernatant samples were collected at 24-, 48-, 72-, and 96-hours post-inoculation and viral titers quantified using One-Step RT-PCR of extracted RNA.

### Competition binning

384-well microtiter plates coated with 1µg/mL of HCoV-HKU1 spike protein were incubated overnight at 4°C. Plates were blocked with 2% BSA in D-PBS for 1 hr at ambient temperature. Purified and unlabeled mAbs, serially diluted tenfold in blocking buffer, were added to the wells (20 µL/well) in quadruplicates and incubated for one hour at ambient temperature. Following incubation, 5 µL of biotin-labeled mAbs at 2.5 µg/mL (a final concentration of 0.5 µg/mL) were added to wells without washing of unlabeled antibodies and incubated for 1hr at ambient temperature. Plates were then washed and developed with HRP-conjugated avidin (Sigma) and a TMB substrate for antibody detection. The signal obtained for binding the biotin-labeled antibody in the presence of an unlabeled tested antibody was expressed as a percentage of the binding of the reference antibody alone after subtracting the background. In this context, ‘competing’ refers to mAbs whose presence reduces the reference antibody binding to less than 70% of its maximum binding, while ‘non-competing’ indicates that the signal was greater than 30%. The levels between 30 to 70% was considered intermediate competition.

### Expression and purification of HKU1-2 Fab for crystallography

The heavy and light chains of the Fab were cloned into the phCMV3 vector. The plasmids were transiently co-transfected into ExpiCHO cells at a ratio of 2:1 (heavy chain to light chain) using ExpiFectamine CHO Reagent (Thermo Fisher Scientific) according to the manufacturer’s instructions. The supernatant was collected at 7 days post-transfection. The Fab was purified with a CaptureSelect CH1-XL Pre-packed Column (Thermo Fisher Scientific), followed by SEC and buffer exchanged 20 mM Tris-HCl, 150 mM NaCl, pH 7.4.

### Ns-EM sample preparation and data collection

Equal concentrations of HCoV-HKU1 spike protein or variants and Fab were diluted to approximately 20 μg/mL with TBS. The sample was directly deposited onto carbon-coated 400-mesh copper grids and stained immediately with 2% (w/v) uranyl formate for 90 seconds. Grids were imaged at 120 KeV on Tecnai T12 Spirit with a 4k × 4k Eagle CCD camera at 52,000× magnification and -1.5 μm nominal defocus. Micrographs were collected using Leginon and the images were transferred to CryoSPARC v4 for processing ^48,49^. Particle stacks were generated in CryoSPARC with particle picking, extraction and iterative rounds of 2D classification and selection. For HCoV-HKU1 spike-mAb complexes, the micrographs were transferred to Relion, where the particles were picked using DoGpicker and classified based on their 2D classes and 3D reconstructions.

### Biolayer interferometry (BLI) binding assays

Binding assays were performed by biolayer interferometry (BLI) using an Octet Red instrument (FortéBio). His8-tagged HCoV-HKU1 spike proteins at 20 μg/mL in 1× kinetics buffer (1x PBS, pH 7.4, 0.01% BSA and 0.002% Tween 20) were loaded onto Ni-NTA sensors, and then incubated with 33 nM, 100 nM, and 300 nM of HKU1-2 IgG. The assay consisted of five steps: 1) baseline: 60 s with 1× kinetics buffer; 2) loading: 360 s with His8-tagged HCoV-HKU1 spike proteins; 3) baseline: 60 s with 1× kinetics buffer; 4) association: 360 s with HKU1-2 IgG; and 5) dissociation: 360 s with 1× kinetics buffer. For estimating the *KD* values, we used a 1:1 binding model.

### Plasmid generation, expression and purification for NTD-NPs and A/Puerto Rico/8/1934 190E hemagglutinin

cDNA encoding HCoV-HKU1 NTD (GenBank: DQ339101; AA 14-294) was cloned into the pcDNA5 expression vector as described previously ^14^. The HCoV-HKU1 NTD was cloned in-frame with a C-terminal human IgG1 Fc, a tobacco etch virus (TEV) cleavage site (ENLYFQG), a 6xHis-tag, and a Strep-tag (WSHPQFEK; IBA, Germany) ^50^. To create the pA-LS NP expression vector ^12^, domain B of protein A (pA; GenBank: M18264.1, AA 212-270) of *Staphylococcus aureus* was fused to 6,7-dimethyl-8-ribityllumazine synthase (LS; GenBank: AAC06489.1, AA 1-154) of *Aquifex aeolicus* by a Gly-Ser linker and to a Strep-tag (WSHPQFEK). The pA-LS sequence was constructed in a pUC57 plasmid by GenScript USA, Inc. and ligated into a pCD5 expression vector via NheI/NotI restriction sites. The A/Puerto Rico/8/1934 190E hemagglutinin protein (GenBank: NP_040980) was cloned into the pCD5 expression vector in-frame with a GCN4 trimerization motif (KQIEDKIEEIESKQKKIENEIARIKK), TEV cleavage site, sfGFP sequence, and Twin-Strep-tag (WSHPQFEKGGGSGGGSGGSAWSHPQFEK; IBA, Germany), as previously described ^36^.

Viral proteins were expressed by transfecting HEK293S GnTI^-^ cells, and pA-LS NP was expressed by transfecting HEK293T cells, as previously described ^51^. Briefly, cells were grown in Dulbecco’s Modified Eagle Medium (DMEM; Gibco) supplemented with 10% fetal calf serum (FCS; Sigma), 25 units/mL of penicillin and 0.025 mg/mL streptomycin (Sigma). Expression vectors were incubated for 20 min. with polyethylenimine I hydrochloride (PEI-HCl) and DMEM in a 1:8 ratio (μg DNA/μg PEI-HCL). The transfection medium was replaced with 293 SFM II medium (Gibco) supplemented with Primatone 3.0 g/L (Kerry), bicarbonate 3.6 g/L, glucose 2.0 g/L, valproic acid 0.4 g/L, glutaMAX 1% (Gibco), and DMSO 1.5% at 6 hours post-transfection. After 5 days of incubation at 37°C and 5% CO2, supernatants were collected, and protein expression was analyzed with SDS-PAGE followed by western blotting using human StrepMAB and goat anti-human IgG HRP (31410, Invitrogen). After purification of proteins with Strep-Tactin Sepharose beads (2-1201-002, IBA) according to the manufacturers protocol, proteins were checked on coomassie blue-stained SDS-PAGE gel.

### Hemagglutination assay

Hemagglutination assays were performed with 100 µL of HCoV-HKU1 NTD NP and A/Puerto Rico/8/1934 190E hemagglutinin, serially diluted with a starting concentration of 20 µg/ml. HCoV-HKU1 NTD was conjugated to pA-LS NP at a 1:1 molar ratio for 1 h on ice, while A/Puerto Rico/8/1934 190E hemagglutinin was incubated with human StrepMAB and goat anti-human IgG HRP (31410, Invitrogen) at a 4:2:1 molar ratio for 30 min on ice. After proteins were 2-fold serially diluted in V-bottom 96-well plates, 50 μL of 1% rat RBCs (*Rattus norvegicus* strain Fischer) were added. HA titers were calculated as the highest dilution of protein hemagglutinating rat RBCs. For depletion of 9-*O*-acetylated Sia, rat RBCs (15% in d-PBS) were treated with 2 µg/mL BCoV HE for 2 h at 37 ⁰C.

### Hemagglutination inhibition assay

Hemagglutination inhibition assay was performed to determine the ability of HKU1-2 mAb to inhibit binding of HCoV-HKU1 NTD to rat RBCs. Serial 2-fold dilutions of each mAb (RSV IgG, HKU1-11 IgG, HKU1-2 IgG, and HKU1-2 Fab) starting at 25 µg/mL, were added onto a V-bottom 96-well plate. The antibodies were incubated with 25 μL of 4 HA units of pA-LS-NTD NPs for 30 min., after which 50 μL of 1% rat RBCs were added. HAI titers were calculated as the highest dilution of mAb capable of hemagglutination inhibition.

### Cryo-EM sample preparation

The HCoV-HKU1 spike protein was incubated with equal molar ratio of HKU1-2 Fab (trimer: Fab = 1:1, final concentration of 0.57 mg/mL) to form spike-HKU1-2 complexes, while spike was complexed with 3-fold molar excess of HKU1-11 Fab (trimer: Fab = 1:3, final concentration of 0.75 mg/mL). The complexes were incubated for 10 and 30 min in TBS (20 mM Tris and 150 mM NaCl, pH 7.4), respectively. For spike-HKU1-11 complexes, 3.5 µL spike-Fab complex was mixed with 0.5 µL of and lauryl maltose neopentyl glycol (LMNG) solution immediately before sample deposition onto an UltrAuFoil 1.2/1.3 (Au, 300-mesh; Quantifoil Micro Tools GmbH) grid that had been plasma cleaned for 25 seconds using a PELCO easiGlow™ Glow Discharge Cleaning System (Ted Pella Inc.). For spike-HKU1-2 complexes, 3.5 µL spike-Fab complex was mixed with 0.5 µL of 0.035 mM n-Octyl-beta-D-glucoside (OBG) prior to sample deposition onto plasma-cleaned Quantifoil 1.2/1.3 (Cu, 300-mesh) grids. Following sample application, grids were blotted for 4 seconds (HKU1-11) and 3 seconds (HKU1-2) before being vitrified in liquid ethane using a Vitrobot Mark IV (Thermo Fisher). The grids were stored in liquid nitrogen before data collection.

### Cryo-EM data collection and processing

Data was collected on Thermo Fisher Glacios, operating at 200 keV mounted with Thermo Fisher Falcon 4 direct electron detector, using the Thermo Fisher EPU 2 software at a magnification of 190,000×. For HKU1-11- and HKU1-2-spike complexes, micrographs were collected at a total cumulative dose of 45.00 e^-^/Å^2^ and 50.14 e^-^/Å^2^ with pixel sizes 0.718 Å and 0.725 Å, respectively. CryoSPARC Live Patch Motion Correction was used for alignment and dose weighting of movies. CTF estimations, particle picking, particle extraction, iterative rounds of 2D classification, ab-initio reconstruction, heterogeneous refinement, homogeneous refinement, 3D Variability, local refinement, and non-uniform refinement were performed on CryoSPARC. The processing workflows for each dataset is shown in **Supplementary Figures S2 and S3**.

### Model building and refinement

Initial model building was performed manually in Coot with HCoV-HKU1 spike (PDB ID: 5I08), and HKU1-2 Fab crystal structure (this study) as templates^31^. HKU1-11 Fab model was predicted by AlphaFold3 ^52^. Followed by iterative rounds of Rosetta relaxed refinement, Phenix real space refinement, and Coot manual refinement to generate the final models ^53-55^. EMRinger and MolProbity were run following each round of Rosetta refinement to evaluate and choose the best refined models ^56,57^. Final map and model statistics are summarized in **Table S1**. Figures were generated using PyMOL, UCSF Chimera, and UCSF Chimera X ^58,59^.

### Crystal structure determination

Purified HKU1-2 Fab (11 mg/mL) was screened for crystallization with the 384 conditions of the JCSG Core Suite (QIAGEN) on our custom-designed robotic CrystalMation system (Rigaku) at Scripps Research by the vapor diffusion method in sitting drops containing 0.1 μL of protein and 0.1 μL of reservoir solution. Diffraction-quality crystals were obtained in 0.04 M potassium dihydrogen phosphate, 20% (v/v) glycerol, and 16% (w/v) polyethylene glycol 8000 at 20°C. Crystals appeared on day 3 and were harvested on day 7. The crystals were then flash-cooled and stored in liquid nitrogen until data collection. Diffraction data were collected at cryogenic temperature (100 K) at National Synchrotron Light Source II (NSLS-II) beamline 17-ID-2 with beam wavelength of 0.97934 Å, respectively. Diffraction data were processed with HKL2000 ^60^. Structures were solved by molecular replacement using PHASER ^61^. Iterative model building and refinement were carried out in COOT ^54^ and PHENIX ^55^, respectively.

### HDX-MS

HCoV-HKU1 spike protein stock solutions (8.3 μM) were prepared in 8.3 mM Tris-HCl, 150 mM NaCl, pH 7.2, and incubated at room temperature (RT) for 30 min. Complexes were formed by incubating HKU1 with a threefold molar excess of HKU1-2 Fab for 30 min at RT. Samples were stored at 0 °C before HDX-MS analysis. Hydrogen/deuterium exchange reactions were initiated by diluting 2 μL of each sample into 4 μL of D₂O buffer (8.3 mM Tris, 150 mM NaCl, pD_read 7.2) and incubating at 0 °C for 10, 100, 1,000, or 10,000 sec. Reactions were quenched with 9 μL of ice-cold buffer (0.1 M glycine, 6.4 M GuHCl, 1 M TCEP, pH 2.4), incubated for 5 min, diluted with 45 μL of 0.1 M glycine (pH 2.4), frozen on dry ice, and stored at -80 °C. Non-deuterated and equilibrium-deuterated controls were prepared as described ^62^. Samples were analyzed using a cryogenic LC system coupled to an Orbitrap Elite mass spectrometer (Thermo Fisher Scientific). Proteolysis was performed online with an immobilized pepsin column (16 μL bed volume, 25 μL/min). Peptides were trapped and separated on an Acclaim PepMap RSLC C18 column (0.3 × 50 mm, 2 μm) with a 5-45% acetonitrile gradient over 30 min. Data were acquired in MS or data-dependent MS/MS mode and analyzed using Proteome Discoverer for peptide identification and HDExaminer (Sierra Analytics) for deuterium uptake calculations with correction for back-exchange ^63-65^.

## References

1 Fehr, A. R. & Perlman, S. Coronaviruses: an overview of their replication and pathogenesis. Methods Mol Biol 1282, 1–23 (2015). 10.1007/978-1-4939-2438-7_1

2 Rabaan, A. A. et al. SARS-CoV-2, SARS-CoV, and MERS-COV: A comparative overview. Infez Med 28, 174–184 (2020).

3 Kesheh, M. M., Hosseini, P., Soltani, S. & Zandi, M. An overview on the seven pathogenic human coronaviruses. Rev Medl Virol 32, e2282 (2022). 10.1002/rmv.2282

4 Liu, D. X., Liang, J. Q. & Fung, T. S. in Encyclopedia of Virology (Fourth Edition) (eds Dennis H. Bamford & Mark Zuckerman) pp428–440 (Academic Press, 2021).

5 Woo, P. C. et al. Characterization and complete genome sequence of a novel coronavirus, coronavirus HKU1, from patients with pneumonia. J Virol 79, 884–895 (2005). 10.1128/jvi.79.2.884-895.2005

6 Lau, S. K. et al. Molecular epidemiology of human coronavirus OC43 reveals evolution of different genotypes over time and recent emergence of a novel genotype due to natural recombination. J Virol 85, 11325–11337 (2011). 10.1128/jvi.05512-11

7 Jackson, C. B., Farzan, M., Chen, B. & Choe, H. Mechanisms of SARS-CoV-2 entry into cells. Nat Rev Mol Cell Biol 23, 3–20 (2022). 10.1038/s41580-021-00418-x

8 Tortorici, M. A. & Veesler, D. Structural insights into coronavirus entry. Adv Virus Res 105, 93–116 (2019). 10.1016/bs.aivir.2019.08.002

9 Walls, A. C. et al. Structure, function, and antigenicity of the SARS-CoV-2 spike glycoprotein. Cell 181, 281–292.e286 (2020). 10.1016/j.cell.2020.02.058

10 Wrapp, D. et al. Cryo-EM structure of the 2019-nCoV spike in the prefusion conformation. Science 367, 1260–1263 (2020). doi:10.1126/science.abb2507

11 Feng, Z. et al. Structural and functional insights into the evolution of SARS-CoV-2 KP.3.1.1 spike protein. Cell Rep 44, 115941 (2025). 10.1016/j.celrep.2025.115941

12 Li, W. et al. Identification of sialic acid-binding function for the middle east respiratory syndrome coronavirus spike glycoprotein. Proc Natl Acad Sci U S A 114, E8508–E8517 (2017). doi:10.1073/pnas.1712592114

13 Hulswit, R. J. G. et al. Human coronaviruses OC43 and HKU1 bind to 9-*O*-acetylated sialic acids via a conserved receptor-binding site in spike protein domain A. Proc Natl Acad Sci U S A 116, 2681–2690 (2019). 10.1073/pnas.1809667116

14 Tomris, I. et al. The HCoV-HKU1 N-terminal domain binds a wide range of 9-*O*-acetylated sialic acids presented on different glycan cores. ACS Infect Dis 10, 3880–3890 (2024). 10.1021/acsinfecdis.4c00488

15 Li, Z. et al. Synthetic *O*-acetylated sialosides facilitate functional receptor identification for human respiratory viruses. Nat Chem 13, 496–503 (2021). 10.1038/s41557-021-00655-9

16 Kim, C. H. SARS-CoV-2 evolutionary adaptation toward host entry and recognition of receptor *O*-acetyl sialylation in virus-host interaction. Int J Mol Sci 21, 4549 (2020). 10.3390/ijms21124549

17 Schultze, B., Gross, H. J., Brossmer, R. & Herrler, G. The S protein of bovine coronavirus is a hemagglutinin recognizing 9-*O*-acetylated sialic acid as a receptor determinant. J Virol 65, 6232–6237 (1991). 10.1128/jvi.65.11.6232-6237.1991

18 Huang, X. et al. Human coronavirus HKU1 spike protein uses *O*-acetylated sialic acid as an attachment receptor determinant and employs hemagglutinin-esterase protein as a receptor-destroying enzyme. J Virol 89, 7202–7213 (2015). doi:10.1128/jvi.00854-15

19 Nguyen, L. et al. Sialic acid-containing glycolipids mediate binding and viral entry of SARS-CoV-2. Nat Chem Biol 18, 81–90 (2022). 10.1038/s41589-021-00924-1

20 Monti, M. et al. Two receptor binding strategy of SARS-CoV-2 is mediated by both the N-terminal and receptor-binding spike domain. J Phys Chem B 128, 451–464 (2024). 10.1021/acs.jpcb.3c06258

21 Tomris, I. et al. SARS-CoV-2 spike N-terminal domain engages 9-*O*-acetylated α2-8-linked sialic acids. ACS Chem. Biol. 18, 1180–1191 (2023). 10.1021/acschembio.3c00066

22 Qing, E., Hantak, M., Perlman, S. & Gallagher, T. Distinct roles for sialoside and protein receptors in coronavirus infection. mBio 11, e02764–02719 (2020). 10.1128/mBio.02764-19

23 Saunders, N. et al. TMPRSS2 is a functional receptor for human coronavirus HKU1. Nature 624, 207-214 (2023). 10.1038/s41586-023-06761-7

24 McCallum, M. et al. Human coronavirus HKU1 recognition of the TMPRSS2 host receptor. Cell 187, 4231–4245. e4213 (2024). 10.1016/j.cell.2024.06.006

25 Fernández, I. et al. Structural basis of TMPRSS2 zymogen activation and recognition by the HKU1 seasonal coronavirus. Cell 187, 4246–4260.e4216 (2024). 10.1016/j.cell.2024.06.007

26 Xia, L., Zhang, Y. & Zhou, Q. Structural basis for the recognition of HCoV-HKU1 by human TMPRSS2. Cell Res 34, 526–529 (2024). 10.1038/s41422-024-00958-9

27 Pronker, M. F. et al. Sialoglycan binding triggers spike opening in a human coronavirus. Nature 624, 201–206 (2023). 10.1038/s41586-023-06599-z

28 Wang, H. et al. TMPRSS2 and glycan receptors synergistically facilitate coronavirus entry. Cell 187, 4261–4271.e4217 (2024). 10.1016/j.cell.2024.06.016

29 Ou, X. et al. Crystal structure of the receptor binding domain of the spike glycoprotein of human betacoronavirus HKU1. Nat Commun 8, 15216 (2017). 10.1038/ncomms15216

30 Bangaru, S. et al. Structural mapping of antibody landscapes to human betacoronavirus spike proteins. Sci Adv 8, eabn2911 (2022). doi:10.1126/sciadv.abn2911

31 Kirchdoerfer, R. N. et al. Pre-fusion structure of a human coronavirus spike protein. Nature 531, 118–121 (2016). 10.1038/nature17200

32 Lieu, R. et al. Rapid and robust antibody Fab fragment crystallization utilizing edge-to-edge beta-sheet packing. PLoS One 15, e0232311 (2020). 10.1371/journal.pone.0232311

33 Jin, M. et al. Human coronavirus HKU1 spike structures reveal the basis for sialoglycan specificity and carbohydrate-promoted conformational changes. Nat Commun 16, 4158 (2025). 10.1038/s41467-025-59137-y

34 Neu, U., Bauer, J. & Stehle, T. Viruses and sialic acids: rules of engagement. Current Opin Struct Biol 21, 610–618 (2011). 10.1016/j.sbi.2011.08.009

35 Park, Y.-J. et al. Structures of MERS-CoV spike glycoprotein in complex with sialoside attachment receptors. Nat Struct Mol Biol 26, 1151–1157 (2019). 10.1038/s41594-019-0334-7

36 Nemanichvili, N. et al. Fluorescent trimeric hemagglutinins reveal multivalent receptor binding properties. J Mol Biol 431, 842–856 (2019). 10.1016/j.jmb.2018.12.014

37 Barnard, K. N. et al. Modified sialic acids on mucus and erythrocytes inhibit influenza A virus hemagglutinin and neuraminidase functions. J Virol 94, e01567–01519 (2020). doi:10.1128/jvi.01567-19

38 Zeng, Q., Langereis, M. A., van Vliet, A. L. W., Huizinga, E. G. & de Groot, R. J. Structure of coronavirus hemagglutinin-esterase offers insight into corona and influenza virus evolution. Proc Natl Acad Sci U S A 105, 9065–9069 (2008). doi:10.1073/pnas.0800502105

39 Gilchuk, I. M. et al. Human antibody recognition of H7N9 influenza virus HA following natural infection. JCI Insight 6, e152403 (2021). 10.1172/jci.insight.152403

40 Díaz-Salinas, M. A., Jain, A., Durham, N. D. & Munro, J. B. Single-molecule imaging reveals allosteric stimulation of SARS-CoV-2 spike receptor binding domain by host sialic acid. Sci Adv 10, eadk4920 (2024). doi:10.1126/sciadv.adk4920

41 Walls, A. C. et al. Unexpected receptor functional mimicry elucidates activation of coronavirus fusion. Cell 176, 1026–1039.e1015 (2019). 10.1016/j.cell.2018.12.028

42 Walls, A. C. et al. Tectonic conformational changes of a coronavirus spike glycoprotein promote membrane fusion. Proc Natl Acad Sci U S A 114, 11157–11162 (2017). doi:10.1073/pnas.1708727114

43 Wang, C. et al. Antigenic structure of the human coronavirus OC43 spike reveals exposed and occluded neutralizing epitopes. Nat Commun 13, 2921 (2022). 10.1038/s41467-022-30658-0

44 Lee, P. S. et al. Receptor mimicry by antibody F045-092 facilitates universal binding to the H3 subtype of influenza virus. Nat Commun 5, 3614 (2014). 10.1038/ncomms4614

45 Jo, G. et al. Structural basis of broad protection against influenza virus by human antibodies targeting the neuraminidase active site via a recurring motif in CDR H3. Nat Commun 16, 7067 (2025). 10.1038/s41467-025-62174-2

46 Petitjean, S. J. L. et al. Multivalent 9-*O*-acetylated-sialic acid glycoclusters as potent inhibitors for SARS-CoV-2 infection. Nat Commun 13, 2564 (2022). 10.1038/s41467-022-30313-8

47 Bangaru, S. et al. A site of vulnerability on the influenza virus hemagglutinin head domain trimer interface. Cell 177, 1136–1152.e1118 (2019). 10.1016/j.cell.2019.04.011

48 Potter, C. S. et al. Leginon: a system for fully automated acquisition of 1000 electron micrographs a day. Ultramicroscopy 77, 153–161 (1999). 10.1016/S0304-3991(99)00043-1

49 Punjani, A., Rubinstein, J. L., Fleet, D. J. & Brubaker, M. A. cryoSPARC: algorithms for rapid unsupervised cryo-EM structure determination. Nat Methods 14, 290–296 (2017). 10.1038/nmeth.4169

50 Rodrigues, E. et al. A versatile soluble siglec scaffold for sensitive and quantitative detection of glycan ligands. Nat Commun 11, 5091 (2020). 10.1038/s41467-020-18907-6

51 de Vries, R. P. et al. The influenza A virus hemagglutinin glycosylation state affects receptor-binding specificity. Virology 403, 17–25 (2010). 10.1016/j.virol.2010.03.047

52 Abramson, J. et al. Accurate structure prediction of biomolecular interactions with AlphaFold 3. Nature 630, 493–500 (2024). 10.1038/s41586-024-07487-w

53 Wang, R. Y.-R. et al. Automated structure refinement of macromolecular assemblies from cryo-EM maps using Rosetta. eLife 5, e17219 (2016). 10.7554/eLife.17219

54 Emsley, P., Lohkamp, B., Scott, W. G. & Cowtan, K. Features and development of Coot. Acta Crystallogr D Biol Crystallogr 66, 486–501 (2010). 10.1107/s0907444910007493

55 Adams, P. D. et al. PHENIX: a comprehensive Python-based system for macromolecular structure solution. Acta Crystallogr D Biol Crystallogr 66, 213–221 (2010). 10.1107/s0907444909052925

56 Williams, C. J. et al. MolProbity: More and better reference data for improved all-atom structure validation. Prot Sci 27, 293–315 (2018). 10.1002/pro.3330

57 Barad, B. A. et al. EMRinger: side chain–directed model and map validation for 3D cryo-electron microscopy. Nat Methods 12, 943–946 (2015). 10.1038/nmeth.3541

58 Goddard, T. D. et al. UCSF ChimeraX: Meeting modern challenges in visualization and analysis. Prot Sci 27, 14–25 (2018). 10.1002/pro.3235

59 Pettersen, E. F. et al. UCSF Chimera—A visualization system for exploratory research and analysis. J Comput Chem 25, 1605–1612 (2004). 10.1002/jcc.20084

60 Otwinowski, Z. & Minor, W. Processing of X-ray diffraction data collected in oscillation mode. Meth Enzymol 276, 307–326 (1997). 10.1016/S0076-6879(97)76066-X

61 McCoy, A. J. et al. Phaser crystallographic software. J. Appl. Crystallogr. 40, 658–674 (2007). 10.1107/S0021889807021206

62 Zost, S. J. et al. Canonical features of human antibodies recognizing the influenza hemagglutinin trimer interface. J Clin Invest 131, e146791 (2021). 10.1172/JCI146791

63 Woods, V. L., Jr. & Hamuro, Y. High resolution, high-throughput amide deuterium exchange-mass spectrometry (DXMS) determination of protein binding site structure and dynamics: utility in pharmaceutical design. J Cell Biochem Suppl **Suppl** 37, 89–98 (2001). 10.1002/jcb.10069

64 Walters, B. T., Ricciuti, A., Mayne, L. & Englander, S. W. Minimizing back exchange in the hydrogen exchange-mass spectrometry experiment. J Am Soc Mass Spectrom 23, 2132–2139 (2012). 10.1007/s13361-012-0476-x

65 Zhang, Z. & Smith, D. L. Determination of amide hydrogen exchange by mass spectrometry: a new tool for protein structure elucidation. Protein Sci 2, 522–531 (1993). 10.1002/pro.5560020404

