## Supplemental Figures and Tables for "Human coronavirus HKU1 neutralization by glycan receptor mimicry"

A

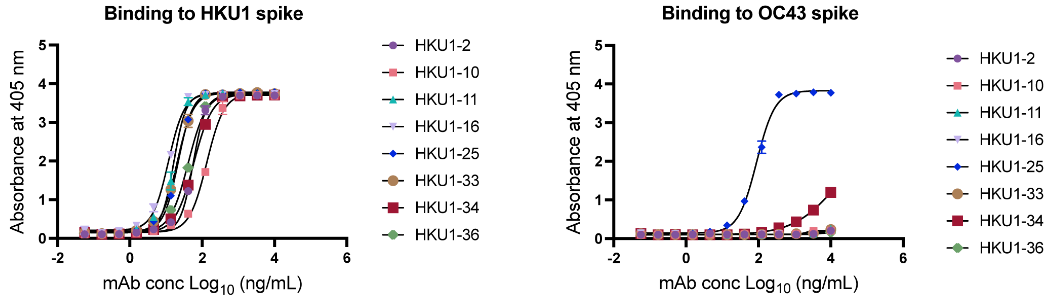


B

**mAb gene usage and CDR3 sequence**

| Clone | Heavy chain V gene | Light chain V gene | HCDR3 | LCDR3 |
|---|---|---|---|---|
| HKU1-2 | IGHV1-24*01 | IGKV3-11*01 | ATDSHCTSRSCYESPREFWYLDV | QQRSNWPPLT |
| HKU1-10 | IGHV1-69*01 | IGKV3-15*01 | AKDLTTVIPYGMDV | QQYNNWPPVYT |
| HKU1-11 | IGHV4-59*08 | IGLV1-47*01 | ARLSCGADCYPIPKLSWFDP | AAWDDSLSGPV |
| HKU1-16 | IGHV1-18*01 | IGKV2-28*01 | ARGNCSTSRCYAVFAY | MQALQTPPT |
| HKU1-25 | IGHV1-69*01 | IGKV3-15*01 | ARVSHAFGGLIVEDWFDP | QQYNNWPPLT |
| HKU1-33 | IGHV3-33*01 | IGKV1-12*01 | GRDKGRYSYGYDAFDF | HQGKSFPRT |
| HKU1-34 | IGHV7-4-1*02 | IGKV3-11*01 | ARAEVMYRGSDY | QHRSNWPPYT |
| HKU1-36 | IGHV3-33*01 | IGKV3-11*01 | ARDRLNWNDPMDV | QQRINWPPLT |

C

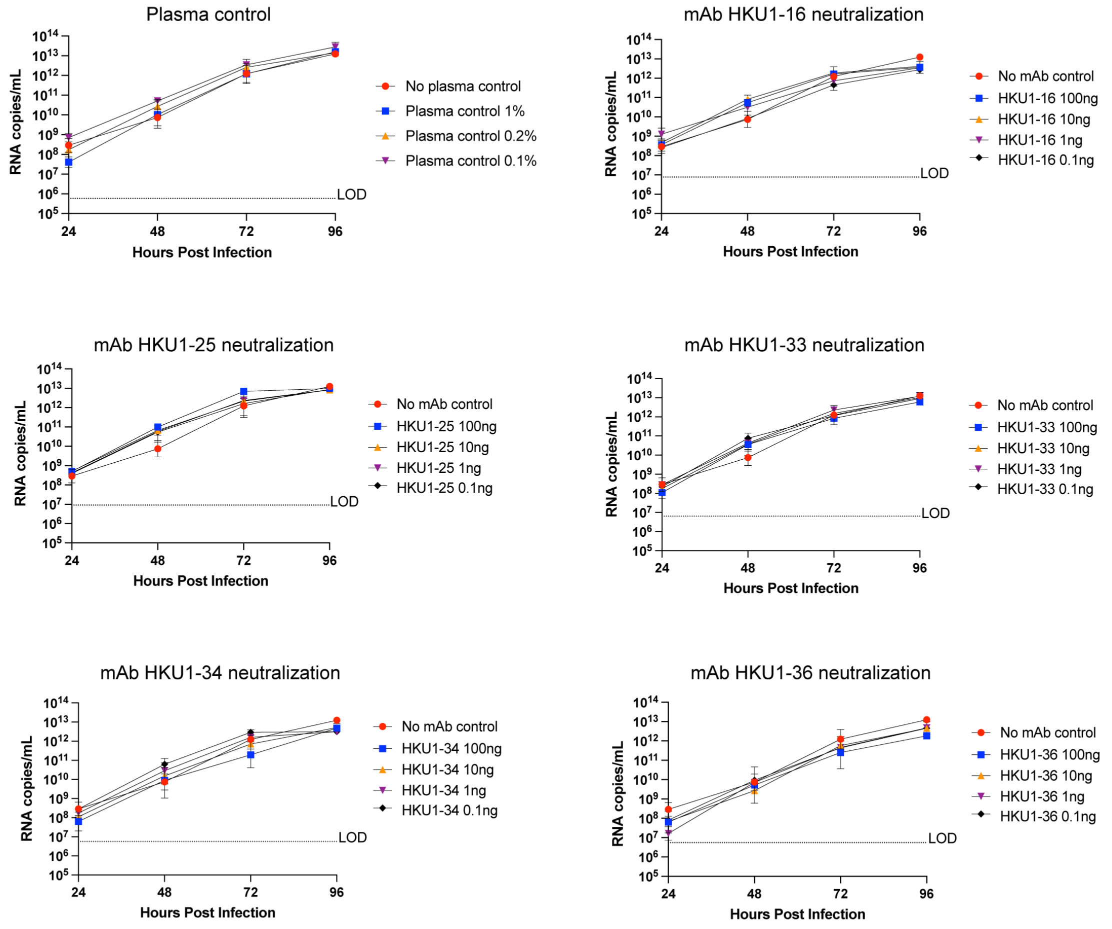

**Figure S1. mAbs neutralization against HCoV-HKU1** (**A**) ELISA binding curves for eight HCoV-HKU1 mAbs to HCoV-HKU1 and HCoV-OC43 spike with a starting concentration of 20 µg/mL. (**B**) Gene usage and CDR3 sequence of eight HKU1 mAbs. (**C**) Inhibition of HCoV-HKU1 viral growth and replication measured at 24, 48, 72 and hours. Replication kinetics of HCoV-HKU1 was assessed by real-time RT-PCR of RNA present in apical washes from HCoV-HKU1-infected human ciliated airway epithelial cells cultures pretreated with either no antibody (red) or with mAb at 100 ng (blue), 10 ng (yellow), 1 ng (violet), or 0.1 ng (black) per mL. Titers are represented by virus RNA copies per milliliter. No significant differences in HCoV-HKU1 viral titers were observed in the antibody treatment groups (two-way ANOVA test). n = 2 for each treatment group.

Schematic representation of the cryo-EM processing workflow in CryoSPARC (HKU1 S + HKU1-11 Fab)

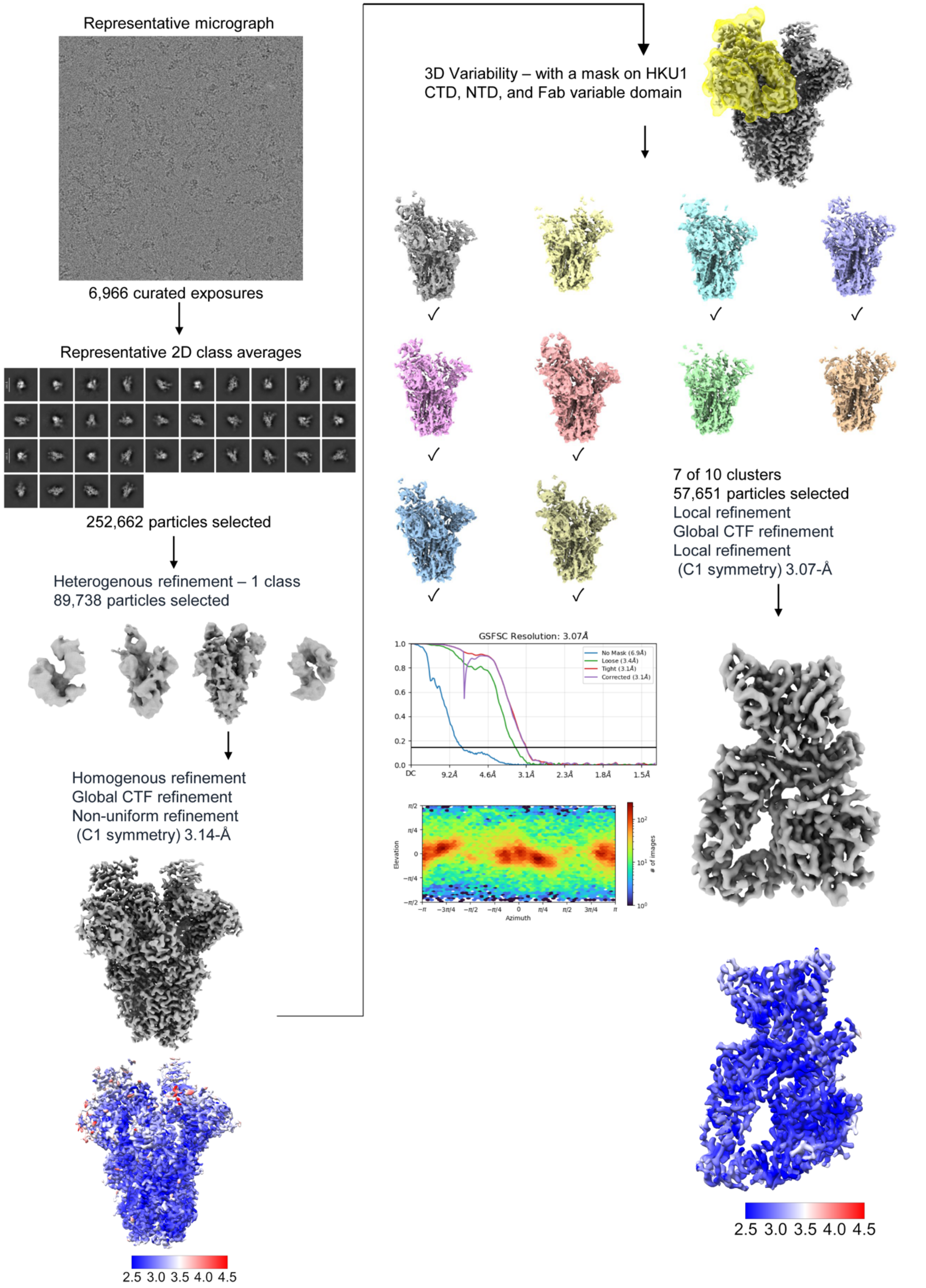

**Figure S2. Schematic representation of the cryo-EM processing workflow for HCoV-HKU1 spike with HKU1-11 Fab.** Cryo-EM processing workflow in cryoSPARC including initial processing steps (motion correction, CTF estimation, micrograph selection, particle picking, and selection based on 2D classification and heterogenous refinement) and 3D variability with a mask on CTD, NTD and Fab. All particles were then separated into 10 clusters and classified by their conformation. 7 of 10 clusters displaying Fab density were selected for further local refinement, which was used for model building. Angular distribution, corresponding FSC curves, and resolution distribution of maps were also displayed.

Schematic representation of the cryo-EM processing workflow in CryoSPARC (HKU1 S + HKU1-2 Fab)

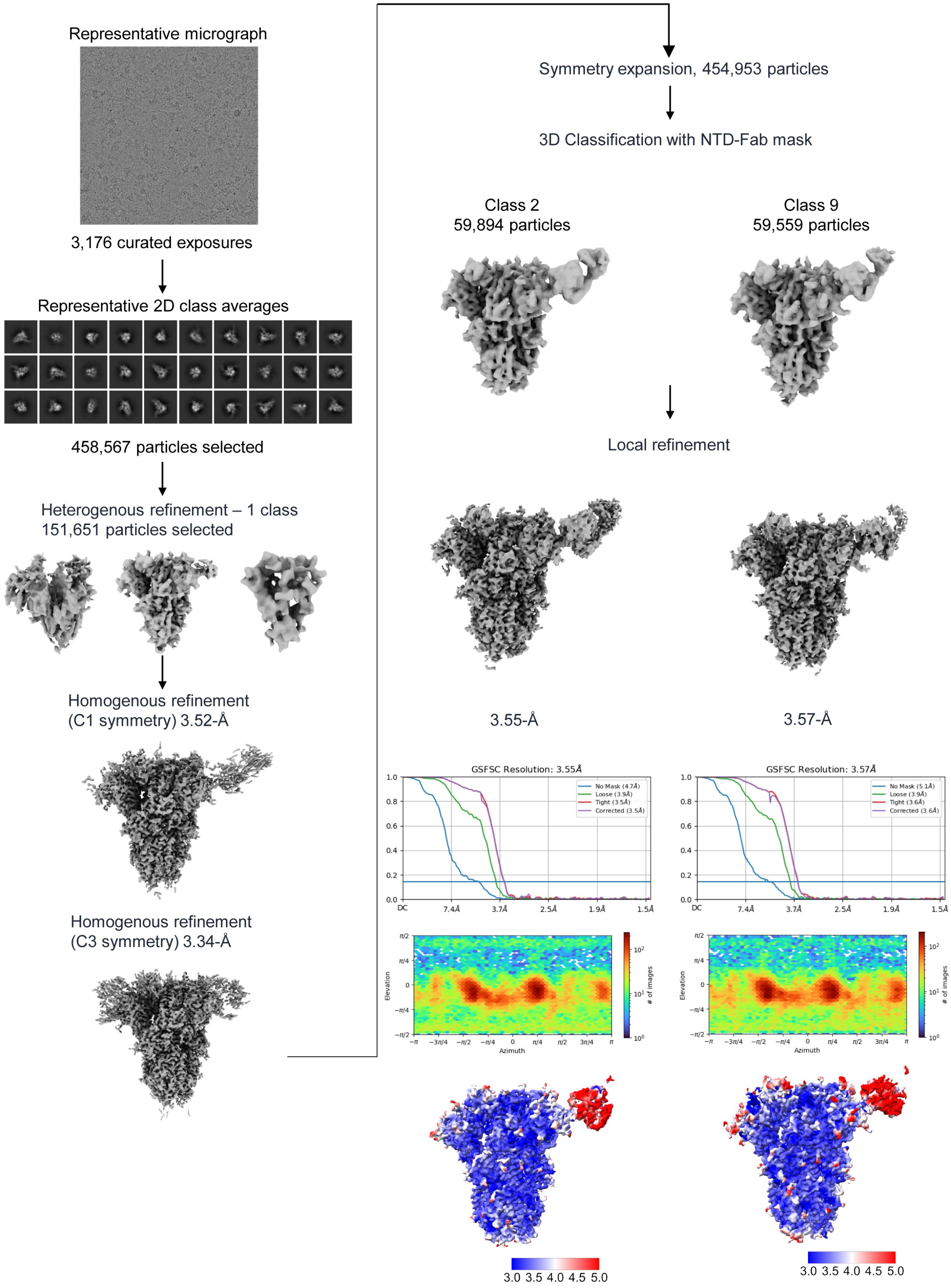

**Figure S3. Schematic representation of the cryo-EM processing workflow for HCoV-HKU1 spike with HKU1-2 Fab.** Cryo-EM processing workflow in cryoSPARC including initial processing steps (motion correction, CTF estimation, micrograph selection, particle picking, and selection based on 2D classification and heterogenous refinement) and 3D classification with a mask on NTD and Fab. All particles were then separated into 10 clusters and classified by their conformation. 2 of 10 clusters displaying Fab density were selected for further local refinement, which was used for model building. Angular distribution, corresponding FSC curves, and resolution distribution were also displayed.

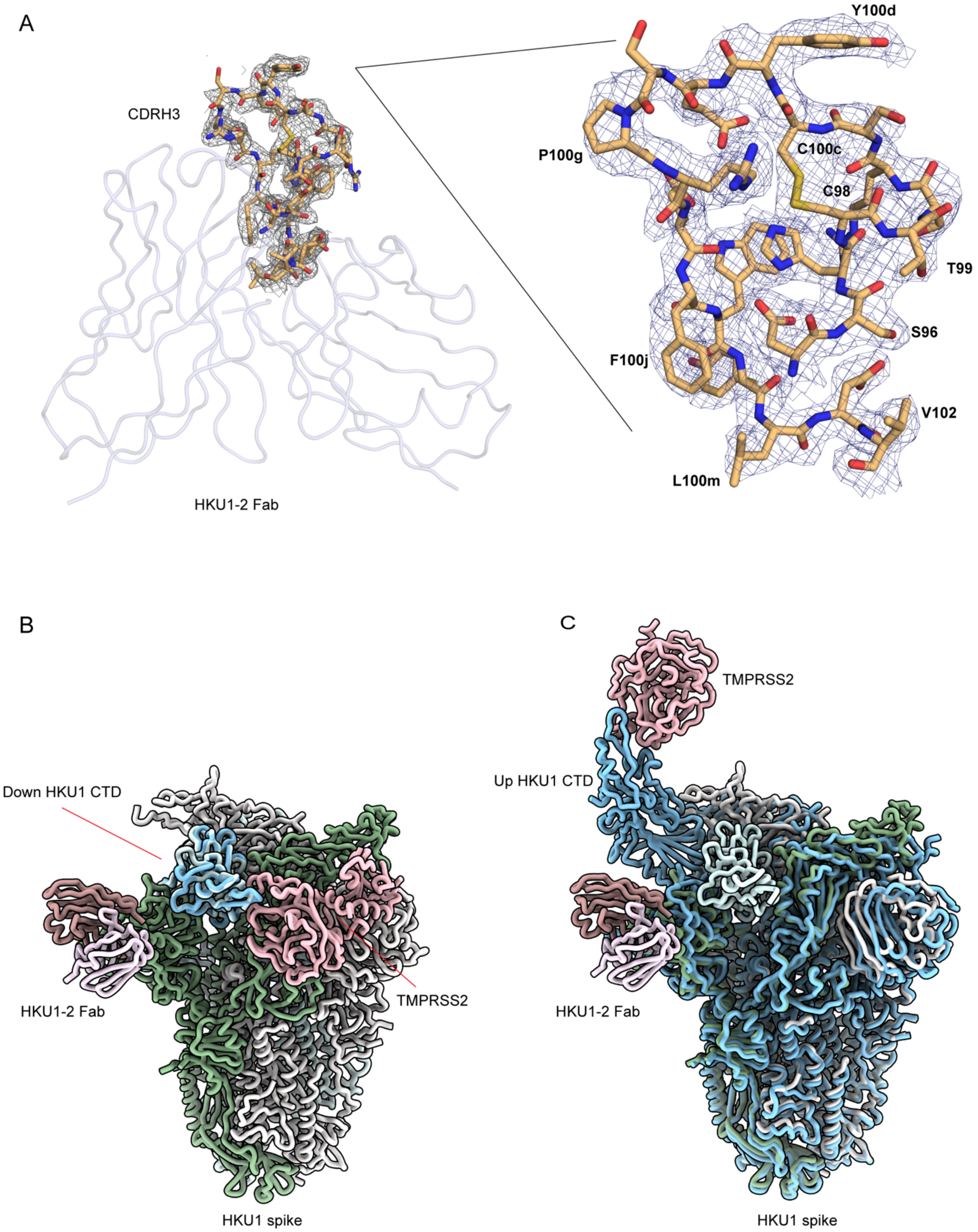


**Figure S4.** (**A**) Electron density maps of HKU1-2 Fab CDRH3 in the crystal structure of apo HKU1-2 Fab. 2Fo-Fc electron density maps for 20 CDRH3 residues are contoured at a 1 σ level and represented in blue mesh with structure showed as sticks. Some residues are labeled. (**B**) Structural superimposition of HCoV-HKU1 CTD/TMPRSS2 (PDB ID: 8VGT) with HCoV-HKU1 spike/HKU1-2 Fab using down HCoV-HKU1 CTD as reference. HCoV-HKU1 spike/HKU1-2 Fab color scheme is same as Figure 3A. HCoV-HKU1 CTD and TMPRSS2 are colored in light blue and pink. TMPRSS2 would not clash with HKU1-2 Fab when the CTD in the down conformation. (**C**) Structural superimposition of HCoV-HKU1 spike/TMPRSS2 (PDB ID: 8Y87) with HCoV-HKU1 spike/HKU1-2 Fab using down HCoV-HKU1 spike as reference. The TMPRSS2-bound CTD is in the up conformation and does not clash with HKU1-2 Fab.

A

```
HKU1-A          -------------VIGDFNCTNFAINDLNTTVPRISEYVVDVSYGLGTYYILDRVYLNTT
HKU1-B          MFLIIFILPTTLAVIGDFNCTNSFINDYNKTIPRISEDVVDVSLGLGTYYVLNRVYLNTT
HKU1-C          -------------VIGDFNCTNSFINDYNKTIPRISEDVVDVSLGLGTYYVLNRVYLNTT
                             *********  *** *.*:***** ***** ******:*:*******

HKU1-A          ILFTGYFPKSGANFRDLSLKGTTYLSTLWYQKPFLSDFNNGIFSRVKNTKLYVNKTLYSE
HKU1-B          LLFTGYFPKSGANFRDLALKGSKYLSTLWYKPPFLSDFNNGIFSKVKNTKLYVNNTLYSE
HKU1-C          LLFTGYFPKSGANFRDLALKGSIYLSTLWYKPPFLSDFNNGIFSKVKNTKLYVNNTLYSE
                :****************:***: *******: ************:*********:*****

HKU1-A          FSTIVIGSVFINNSYTIVVQPHNGVLEITACQYTMCEYPHTICKSKGSSRNESWHFDKSE
HKU1-B          FSTIVIGSVFVNTSYTIVVQPHNGILEITACQYTMCEYPHTVCKSKGSIRNESWHIDSSE
HKU1-C          FSTIVIGSVFVNTSYTIVVQPHNGILEITACQYTMCEYPHTVCKSKGSIRNESWHIDSSE
                **********:*.***********:****************:****** ******:*.**

HKU1-A          PLCLFKKNFTYNVSTDWLYFHFYQERGTFYAYYADSGMPTTFLFSLYLGTLLSHYYVLPL
HKU1-B          PLCLFKKNFTYNVSADWLYFHFYQERGVFYAYYADVGMPTTFLFSLYLGTILSHYYVMPL
HKU1-C          PLCLFKKNFTYNVSADWLYFHFYQERGVFYAYYADVGMPTTFLFSLYLGTILSHYYVMPL
                **************:************.******* **************:******:**

HKU1-A          TCNAISSNTDNETLQYWVTPLSKRQYLLKFDNRGVITNAVDCSSSFFSEIQCKTKSLLPN
HKU1-B          TCKAISSNTDNETLEYWVTPLSRRQYLLNFDEHGVITNAVDCSSSFLSEIQCKTQSFAPN
HKU1-C          TCNAISSNTDNETLEYWVTPLSRRQYLLNFDEHGVITNAVDCSSSFLSEIQCKTQSFAPN
                **:***********:*******:*****:**::*************:*******:*: **
```

B

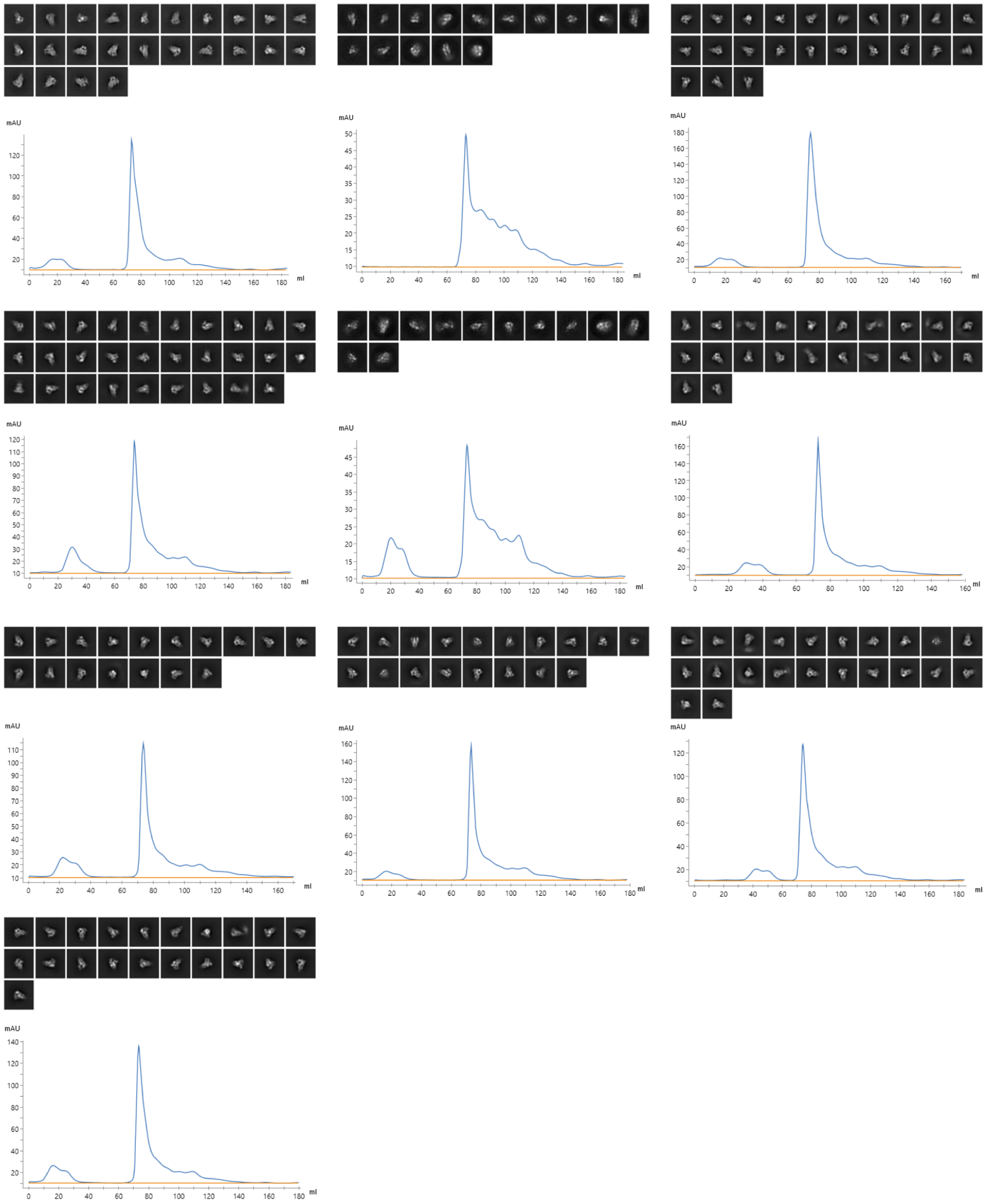

**Figure S5.** (**A**) Sequence alignment of HCoV-HKU1 NTD of three genotypes A, B, and C. Yellow highlights six conserved epitope residues which also participate in the antibody interactions. Residues identical in all aligned sequences are labeled by an asterisk (*), whereas a colon (:) and a period (.) indicate sequences assigned as strongly similar and less similar, respectively[1]. The sequence alignment was performed with Clustal Omega[1]**.** (**B**) Representative 2D class averages and size-exclusion chromatography traces of various HCoV-HKU1 spike mutants.

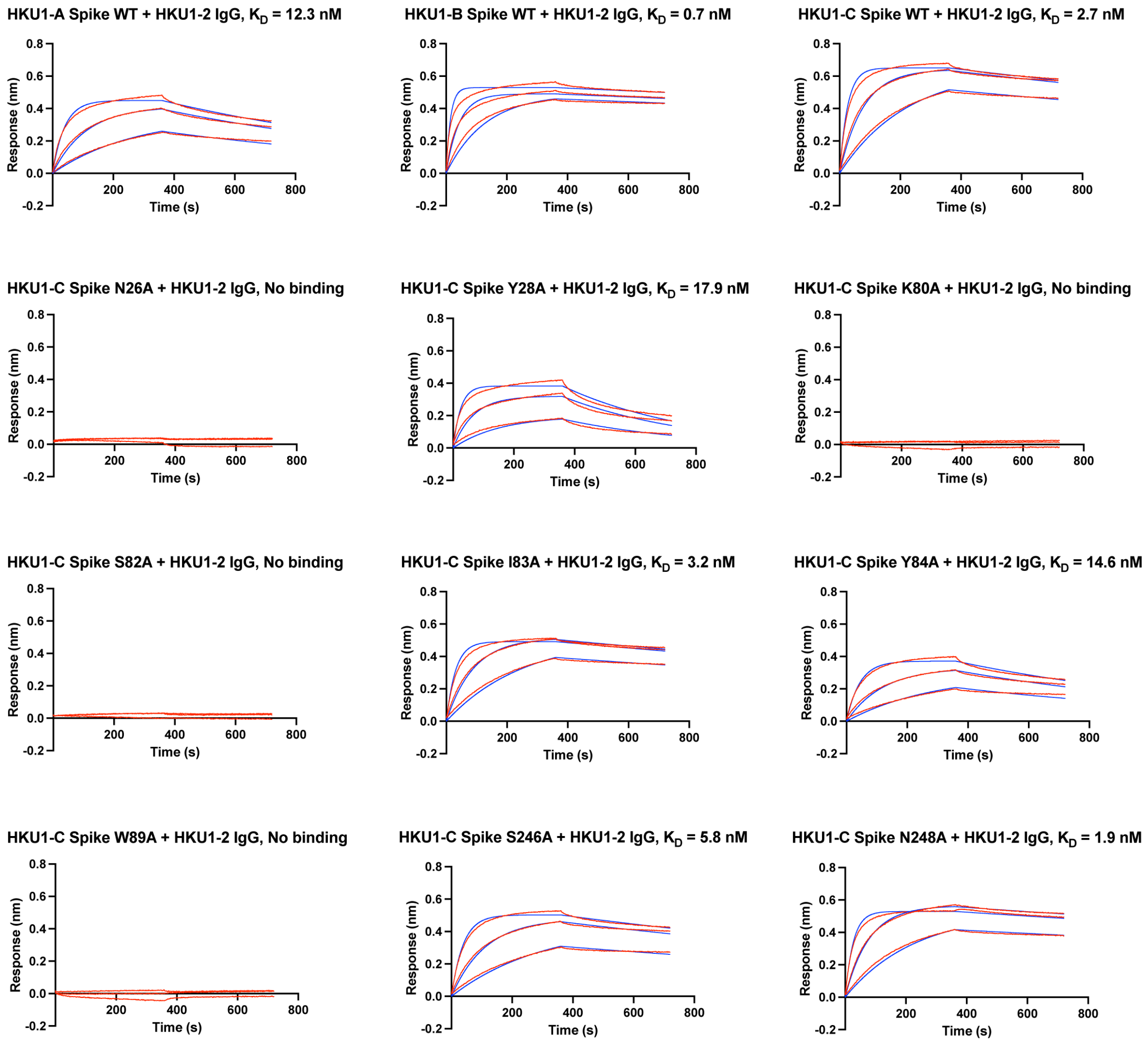


**Figure S6. Sensorgrams for binding of various HCoV-HKU1 spike proteins and mutants to HKU1-2 IgG.** Binding kinetics for HKU1-2 IgG to spikes derived from different HCoV-HKU1 genotypes and mutants measured by biolayer interferometry (BLI). The Y-axis represents the response. Red lines represent the response curve, and blue lines represent a 1:1 binding model. Binding kinetics were measured for HKU1-2 IgG concentrations (33.3, 100, and 300). Binding affinity is expressed as the nanomolar dissociation constant ($K_D$).

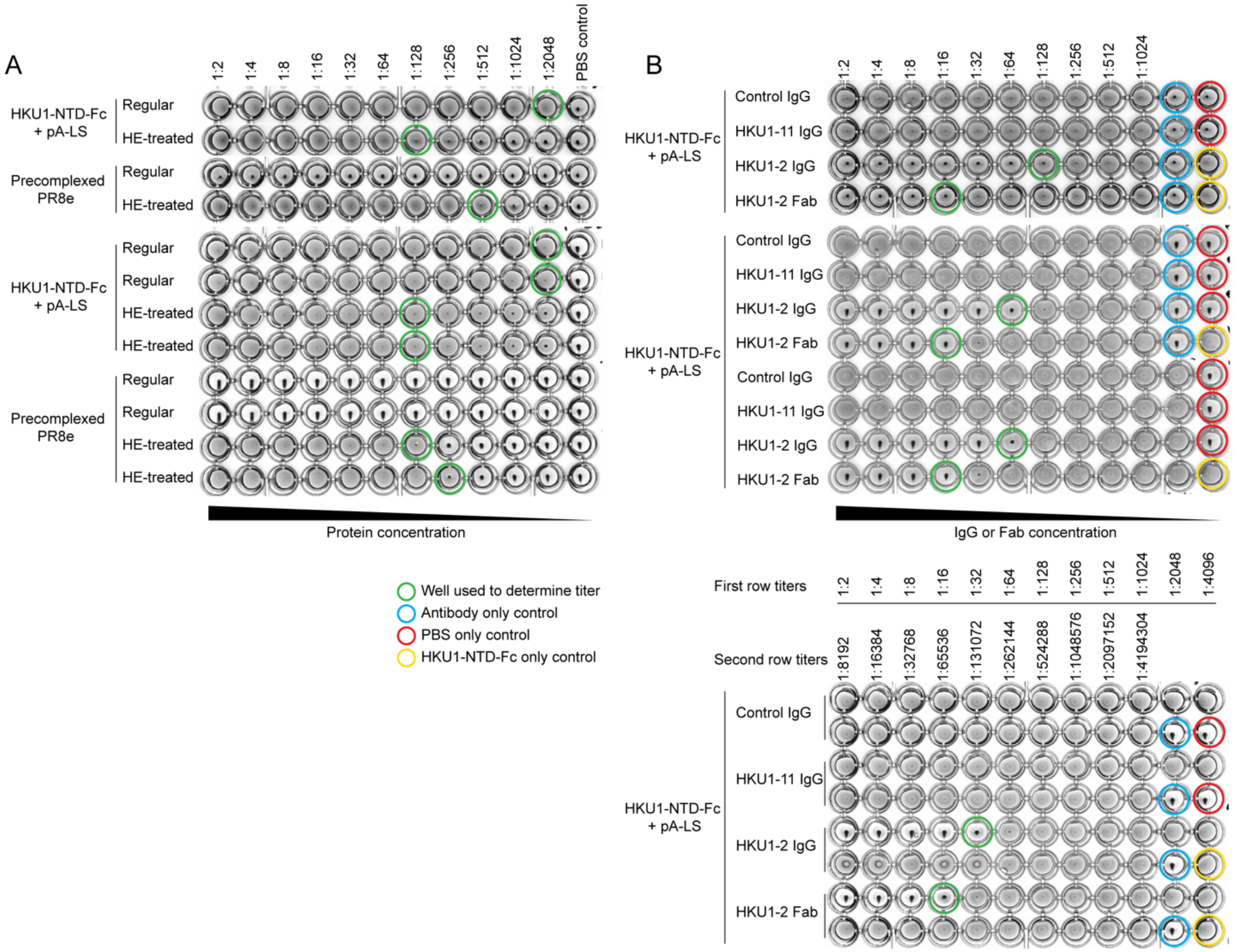


**Figure S7 HKU1-2 mediated inhibition of HKU1 hemagglutination of rat RBCs** (**A**) Plates for hemagglutination assay. Hemagglutination of rat RBCs by HCoV-HKU1 NTD NP and influenza A/Puerto Rico/8/1934 HA was assessed using two-fold serial dilution in a 96-well V-bottom plate. For depletion of 9-*O*-acetylated sialic acid, BCoV HE was added. Green circle highlights the wells used for titer determination. 4 HA units was used to perform hemagglutination inhibition assay (**B**) Plates for hemagglutination inhibition assay. Hemagglutination inhibition was assessed by two-fold serial dilution of mAbs (RSV IgG, HKU1-11 IgG, HKU1-2 IgG, and HKU1-2 Fab) in a 96-well V-bottom plate. Green circle highlights the wells used for titer determination. Blue, red, and yellow circles represent antibody, PBS, and HKU1-NTD only control, respectively.

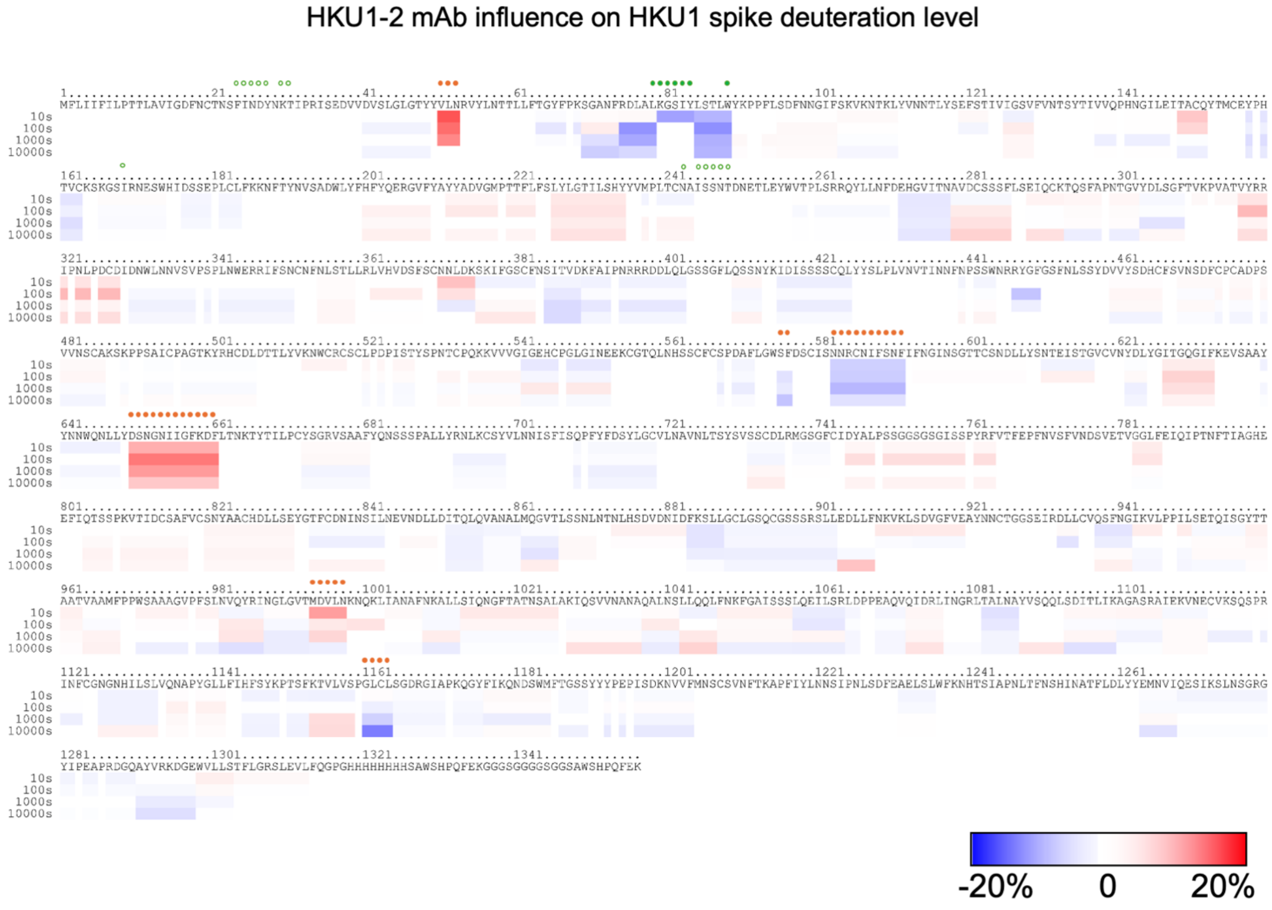


**Figure S8. HDX-MS reveals changes in deuteration levels for HKU1-2 mAb bound spike compared to apo HCoV-HKU1 spike protein.** The differences in deuteration are indicated on the spike sequence as a color gradient: blue for decrease, red for increase of deuteration levels. Green circles indicate amino acid residues within the epitope, as determined by cryo-EM. Solid green circles indicate amino acid residues with decreased deuteration level within the epitope upon mAb HKU1-2 binding to HCoV-HKU1 spike protein. Solid orange circles indicate amino acid residues with increased or decreased deuteration level outside of the HKU1-2 binding epitope. Each row indicates the deuteration differences at different time points.

**Table S1. Cryo-EM data collection, processing, and model building statistics.**

| Sample | HKU1 Spike (S2P) + HKU1-2 Fab | | HKU1 Spike (S2P) + HKU1-11 Fab |
|---|---|---|---|
| **Map** | State 1 | State 2 | Local refinement |
| **EMDB** | EMD-73659 | EMD-73660 | EMD-73658 |
| **PDB** | 9YYY | 9YYZ | 9YYX |
| **Data collection & processing** | | | |
| **Microscope/ Detector** | Glacios/Falcon4 | | |
| **Voltage (kV)** | 200 | | |
| **Magnification** | 190,000 | | |
| **Recording mode** | Counting | | |
| **Pixel size (Å)** | 0.718 | | |
| **Total dose (e-/Å$^2$)** | 50.14 | | 45.00 |
| **Defocus range (μm)** | -0.7 to -1.7 | | |
| **No. of movie micrographs** | 3,176 | | 6,966 |
| **No. of molecular projection images in map** | 59,894 | 59,559 | 57,651 |
| **Symmetry** | C1 | C1 | C1 |
| **Map pixel size (Å)** | 0.725 | | 0.718 |
| **Map resolution (FSC 0.143; Å)** | 3.55 | 3.57 | 3.07 |
| **Map sharpening B-factor (Å$^2$)** | -55.0 | -53.0 | -71.5 |
| **Structure building and validation** | | | |
| **Model composition** | | | |
| **Non-hydrogen atoms** | 31,113 | 31,131 | 6,537 |
| **Protein residues** | 3,813 | 3,813 | 810 |
| **Ligands** | NAG: 82/ BMA: 11/ MAN: 6 | NAG: 82/ BMA: 11/ MAN: 6 | NAG: 12/ BMA: 2/ FUC: 1 |
| **RMSD bond length (Å)/angles (°)** | 0.007/1.216 | 0.007/1.139 | 0.008/1.299 |
| **MolProbity score** | 1.14 | 0.91 | 1.22 |
| **Clash score** | 1.71 | 1.03 | 1.33 |
| **Ramachandran outliers/allowed/favored (%)** | 0.00/3.36/96.64 | 0.00/2.51/97.49 | 0.00/5.12/94.88 |
| **Rotamer outliers (%)** | 0.00 | 0.03 | 0.00 |
| **Cβ outliers (%)** | 0.00 | 0.03 | 0.13 |
| **d FSC model (0.5; Å)** | 3.9 | 3.9 | 3.3 |

### Table S2. X-ray data collection and refinement statistics

| Data collection | HKU1-2 Fab |
|---|---|
| Beamline | NSLS-II 17-ID-2 |
| Wavelength (Å) | 0.97934 |
| Space group | $P\ 2_1\ 2_1\ 2_1$ |
| Unit cell parameters | |
| a, b, c (Å) | 48.7, 74.0, 145.9 |
| α, β, γ (°) | 90.0, 90.0, 90.0 |
| Resolution (Å) [a] | 50.0-2.14 (2.18-2.14) |
| Unique reflections [a] | 30,045 (1,357) |
| Redundancy [a] | 7.2 (4.0) |
| Completeness (%) [a] | 99.1 (90.5) |
| $<I/\sigma_I>$ [a] | 17.8 (1.8) |
| $R_{sym}$ [b] (%) [a] | 9.9 (59.3) |
| $R_{pim}$ [b] (%) [a] | 3.8 (31.7) |
| $CC_{1/2}$ [c] (%) [a] | 99.7 (59.9) |
| **Refinement statistics** | |
| Resolution (Å) | 46.2-2.14 |
| Reflections (work) | 28,286 |
| Reflections (test) | 1,987 |
| $R_{cryst}$ [d] / $R_{free}$ [e] (%) | 18.8/23.6 |
| Copies of Fab per ASU | 1 |
| No. of atoms | 3,820 |
| Fab | 3,396 |
| Ligands [f] | 18 |
| Solvent | 406 |
| Average $B$-values (Å$^2$) | 32 |
| Fab | 32 |
| Ligands [f] | 36 |
| Solvent | 34 |
| Wilson $B$-value (Å$^2$) | 27 |
| **RMSD from ideal geometry** | |
| Bond length (Å) | 0.002 |
| Bond angle (°) | 0.60 |
| **Ramachandran statistics (%)** [g] | |
| Favored | 98.6 |
| Outliers | 0.0 |
| **PDB code** | 9YYW |

[a] Numbers in parentheses refer to the highest resolution shell.

[b] $R_{sym} = \Sigma_{hkl} \Sigma_i | I_{hkl,i} - <I_{hkl}> | / \Sigma_{hkl} \Sigma_i I_{hkl,i}$ and $R_{pim} = \Sigma_{hkl} (1/(n-1))^{1/2} \Sigma_i | I_{hkl,i} - <I_{hkl}> | / \Sigma_{hkl} \Sigma_i I_{hkl,i}$, where $I_{hkl,i}$ is the scaled intensity of the i$^{th}$ measurement of reflection h, k, l, $<I_{hkl}>$ is the average intensity for that reflection, and $n$ is the redundancy.

[c] $CC_{1/2}$ = Pearson correlation coefficient between two random half datasets.

[d] $R_{cryst} = \Sigma_{hkl} | F_o - F_c | / \Sigma_{hkl} | F_o |$ x 100, where $F_o$ and $F_c$ are the observed and calculated structure factors, respectively.

[e] $R_{free}$ was calculated as for $R_{cryst}$, but on a test set comprising 7% of the data excluded from refinement.

[f] Bound ligand is the glycerol molecule.

[g] From MolProbity [2].

Reference:

1 Sievers, F. *et al.* Fast, scalable generation of high-quality protein multiple sequence alignments using Clustal Omega. *Mol Syst Biol* **7**, 539 (2011). https://doi.org/10.1038/msb.2011.75

2 Chen, V. B. *et al.* MolProbity: all-atom structure validation for macromolecular crystallography. *Acta Crystallogr. D Biol. Crystallogr.* **66**, 12-21 (2010). https://doi.org/10.1107/S0907444909042073
